# Different dopaminergic circuits defined by D2/D3 receptor availablity patterns show structure-specific links to memory in ageing

**DOI:** 10.1101/2025.07.17.665393

**Authors:** Yeo-Jin Yi, Jarkko Johansson, Berta Garcia-Garcia, Peter Schulze, Kathrin Baldauf, Ines Bodewald, Marianne Patt, Swen Hesse, Andeas Schildan, Philipp Genseke, Simone Schenke, Matthew J. Betts, Joel Dunn, Christin Y. Sander, Osama Sabri, Michael C. Kreissl, Oliver Speck, Emrah Düzel, Dorothea Hämmerer

## Abstract

Cognitive ageing is marked by progressive decline in episodic memory and dopaminergic function, yet the extent to which individual differences in dopaminergic system integrity influence memory under motivational contexts remains unclear. In this study, we investigated how baseline D2/D3 receptor availability (BP_ND_^0^) in key dopaminergic pathways relates to reward-modulated memory performance in healthy older adults. Thirty-three healthy seniors (aged 64-85) underwent two session concurrent MR-PET imaging with [^1^F]fallypride involving scene categorisation task with high- and low-motivational contexts. We quantified BP_ND_^0^ across nine dopaminergic regions of interest and examined their relationships with recognition memory performance at short (∼15m) and long (∼24h) delays.

Baseline D2/D3 receptor availability showed high test-retest reliability and regionally distinct profiles, suggesting it reflects a stable neurochemical characteristic in healthy ageing, and principal axis factor analysis revealed two partially independent dopaminergic subsystems (dorsal striatal vs mesolimbic) based on interindividual patterns in receptor densities. Region-specific associations further linked D2/D3 receptor availability to distinct memory outcomes. Higher caudate D2/D3 receptor availability was associated with a liberal response bias (increased hits and false alarms both), whereas higher putamen D2/D3 predicted more durable long-term memory retention. Greater thalamic D2/D3 receptor availability correlated with fewer short-term false memories, while greater D2/D3 receptor availability in the amygdala was associated with better recognition at longer delays. In contrast, higher midbrain (substantia nigra and ventral tegmental area) D2/D3 availability which was linked to poorer reward-related memory performance. These findings suggest several complementary dopaminergic circuits supporting episodic memory. Our results highlight dopaminergic neuromodulation as a key factor in cognitive ageing and a potential target for interventions to bolster memory in late life.

**Patient consent and ethics approval:** All subjects who were included in this study have provided their consent prior to their participation, following the guidelines of the ethics committee at University Hospital Magdeburg in Magdeburg, Germany.

## Introduction

Ageing is characterised by progressive structural atrophy and neurochemical changes in the brain, with episodic memory exhibiting faster rate of decline compared to other memory types ^1–3^. The neurobiological substrates of this deterioration include grey matter loss, reduced white matter integrity, and compromised neuromodulatory systems ^4–6^. Among these, the dopaminergic system, critical for learning, reward, and episodic memory, undergoes notable age-related changes, including declining dopamine (DA) receptor availability and neurotransmission efficiency ^7–9^. Such declines are strongly associated with reduced cognitive flexibility, processing speed, and memory performance ^10–14^.

In young adults, motivational salience and reward anticipation robustly facilitates episodic memory encoding and consolidation ^15–17^. Reward-associated cues not only strengthen memory for target items but can also bolster the retention of temporally proximate information ^18,19^. This phenomenon is closely linked to dopaminergic activity, which modulates synaptic plasticity underlying memory consolidation ^16,20,21^. However, because relevant observations in older adults is mostly cross-sectional, it remains unclear whether those with relatively preserved dopaminergic function in fact translate salience signals into better episodic memory, or whether observed links simply reflect inter-individual variability without implying resilience or susceptibility. Ageing may alter the link between dopaminergic modulation and memory processes, limiting the efficacy of reward-based strategies to enhance memory performance ^22–24^.

To address these gaps, our study employed concurrent MR-PET imaging with [^18^F]fallypride, a high-affinity D2/D3 receptor antagonist, to quantify inter-individual variation in D2/D3 receptor availability (represented as baseline binding potential [BP_ND_^0^]) across subcortical regions in healthy older adults and to examine (a) the test-retest reliability of these measures across two scan sessions; (b) whether ageing is associated with coordinated regional differences in receptor availability; (c) the correspondence between receptor availability and local grey-matter volume; and (d) how regional receptor availability relates to cognitive components that support reward-associated episodic memory. By focusing on BP_ND_^0^, a proxy measure of dopaminergic system integrity in each region of interest (ROI), and volumetric information of those ROIs, we aim to delineate how interindividual differences in the integrity of the dopaminergic system are linked to episodic encoding, consolidation, and retrieval in an ageing sample. Elucidating the more direct relationship between the integrity of dopaminergic neuromodulation, brain structure, and episodic memory function has practical implications for interventions aimed at preventing or slowing memory loss, which is a hallmark symptoms of various neurodegenerative diseases^12,25^ as well as memory decline observed in normal ageing^1,26^. By identifying links between interindividual differences in D2/D3 receptor availability in specific brain regions and components of episodic memory, we can clarify how age-related declines in dopaminergic function might contribute to age-related decline in memory.

Specifically, we hypothesised that baseline dopaminergic receptor availability in key source regions (e.g. the substantia nigra [SN], ventral tegmental area [VTA]) and target regions (e.g. the hippocampus, amygdala, thalamus, striatum [caudate and putamen]) would predict interindividual differences in reward-associated memory performance. Moreover, we anticipated that reward salience would elicit better memory performance, and higher BP_ND_^0^ in DA-rich areas would relate to stronger recognition performance, particularly for reward-associated stimuli, reflecting a more favourable neuromodulatory environment for encoding. In addition, because structural brain deterioration and dopaminergic dysfunction often co-occur in older adults, we examined volumetric measures normalised with total intracranial volume in those same regions, predicting that relative volumetric preservation might be related to lower receptor densities, thereby better memory outcomes. Finally, by clustering dopaminergic regions based on shared patterns of decline across individuals, we sought to identify subgroups or subnetworks within the dopaminergic system that may differentially contribute to age-related cognitive decline.

## Results

### Baseline Dopaminergic Receptor Availability (BP_ND_^0^)

PET measures of baseline D2/D3 receptor availability (BP_ND_^0^) were examined with a two-factor repeated measures ANOVA (Reward Context [high vs. low] × Region [9 ROIs]; Figure 1). No main effect of Reward Context emerged, *F*(1,32)=0.071, *p*=0.791, confirming stable BP ^0^ across the two within-person test sessions regardless of anticipated reward magnitude. There was, however, a strong main effect of Region, *F*(8,236)=2236.147, *p*<0.001. Striatal regions (caudate, putamen) showed the highest BP_ND_^0^ values, whereas regions such as the amygdala, hippocampus, LC, SN, VTA, and thalamus exhibited relatively lower values. This distribution matches known receptor densities and dopaminergic innervation patterns^27–31^.

**Figure 1.**
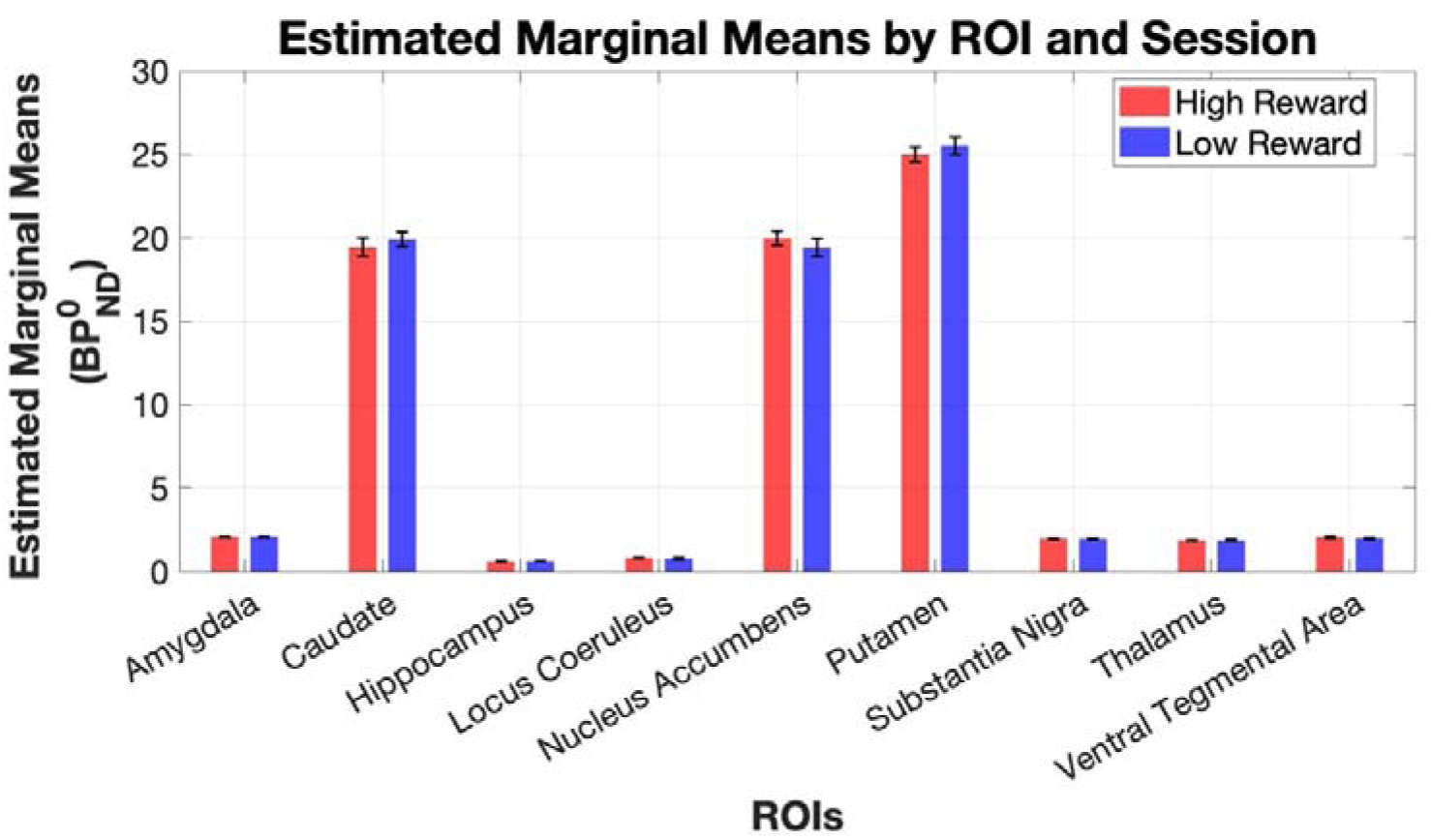
BP_ND_^0^ across dopaminergic ROIs in high- and low-reward contexts. **Data are shown for** both high-reward (red bars) and low reward (blue bars) context test sessions. Error bars represent the standard error of the mean.

### Interindividual Stability and Inter-Regional Relationships of Baseline Receptor Availability

Examination of cross-session relationships revealed that BP_ND_^0^ is highly reproducible across the six-week interval for most regions, i.e. amygdala, caudate, hippocampus, thalamus, SN, VTA, and LC. Prior test-retest studies with [¹_F]fallypride in both young and older adults show intra-class correlations of ≥0.80 and variability below 8% across 1- to 6-week intervals ^32–34^. Consistent with those reports, our data indicate that estimated D2/D3 receptor binding potential is highly stable over weeks, implying that appreciable changes in receptor availability would be expected only over much longer periods (Figure 2A). The striatal BP_ND_^0^, however, was not uniform in our data. The nucleus accumbens (NAcc) did not show a similar cross-session association, and the putamen showed only a trend-level correlation, ρ(31)=0.298, *p*=0.093. These differences might be best explained by region-specific measurement idiosyncrasies rather than a rapid biological change. The NAcc is a relatively small structure embedded between high-uptake structures such as ventral caudate and anterior putamen and low-uptake structures such as prefrontal cortex. Therefore, partial-volume leaks from these regions may significantly influence interscan reliability. Indeed, several D2/D3 receptor PET test-retest studies report intraclass correlation coefficients of ≤0.60 for the NAcc compared to ≥0.80 in dorsal striatum ^33,35,36^. In the putamen, its dense DA innervation might have rendered it especially sensitive to transient endogenous fluctuations from spontaneous movements during this study’s 6-hour visit, which may have suppressed interscan BP_ND_ ^0^ correlations^37^. Moreover, in the putamen, its steep rostrocaudal receptor gradient^38^ (<20-25%) may have reduced observable between-subject variance when averaged across^39^. Collectively, the present result still supports the view that BP_ND_^0^ provides a stable, charicteristic index of dopaminergic system integrity across differing motivational contexts.

**Figure 2.**
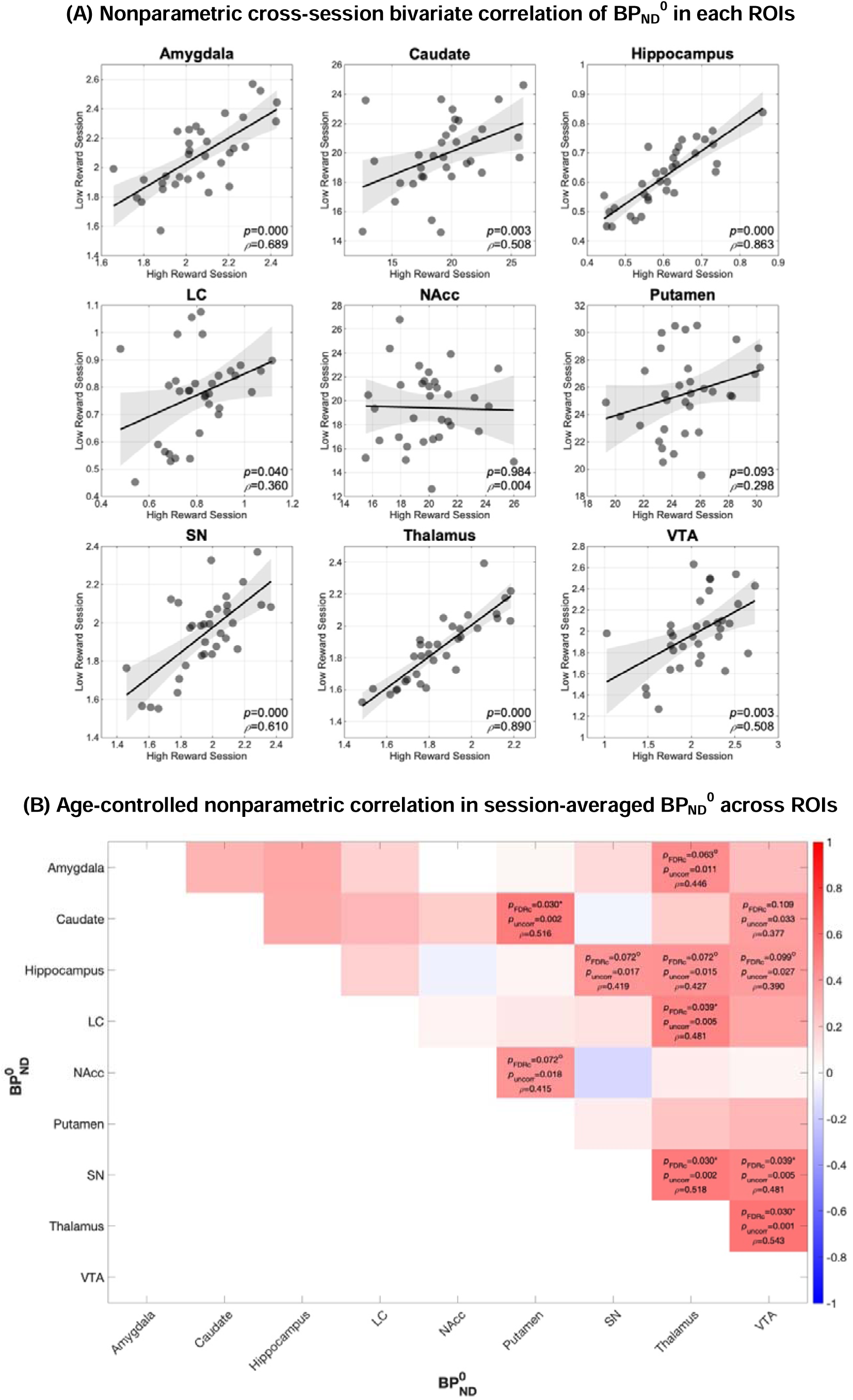
Cross-Session and ROI-averaged nonparametric (Spearman) correlations of baseline D2/D3 receptor availability (BP_ND_^0^). **(A)** Cross-session Spearman’s *bivariate* correlation scatterplots showing the relationships between BP_ND_^0^ of the high-reward session (x-axis) and the low-reward session (y-axis). (**B)** Partial correlation matrix showing cross ROI relationships in session-averaged BP_ND_^0^, age as a control variable. The colour scale indicates Spearman’s ρ values, ranging from −1 (blue) to +1 (red). Each cell displays the correlation coefficient (ρ) and FDR-adjusted p-value (*p*_FDRc_). P-values with asterisk superscripts (*) denote correlations that reached significance of *p*_FDRc_ <.05 and circle superscripts (°) denote *p*_FDRc_<0.1.

Relationships across all dopaminergic ROIs were also assessed with age-controlled, non-parametric partial correlations after averaging BP_ND_^0^ of both sessions per subject to examine the intrinsic organisation of the dopaminergic system in healthy older adults (Figure 2B). Among striatal regions, the caudate and putamen showed the strongest association, ρ(31)=0.516, *p*_uncorr_=0.002, *p*_FDRc_=0.030, while the NAcc moderately correlated with the putamen, ρ(31)=0.415, *p*_uncorr_=0.018, *p*_FDRc_=0.072. A limbic-midbrain covariance was also evident in the result. Hippocampal BP ^0^ showed positive correlation with both SN, ρ(31)=0.419, *p*_uncorr_=0.017, *p*_FDRc_=0.072, and VTA, ρ(31)=0.390, *p*_uncorr_=0.027, *p*_FDRc_=0.099. The thalamus showed multiple correlations with broad range of dopaminergic ROIs, relating not only to limbic structures, hippocampus: ρ(31)=0.427, *p*_uncorr_=0.015, *p*_FDRc_=0.072; amygdala: ρ(31)=0.446, *p*_uncorr_=0.011, *p*_FDRc_=0.063, but also to neuromodulatory nuclei in the brainstem, LC: ρ(31)=0.481, *p*_uncorr_=0.005, *p*_FDRc_=0.039; SN: ρ(31)=0.518, *p*_uncorr_=0.002, *p*_FDRc_=0.030; VTA: ρ(31)=0.543, *p*_uncorr_=0.001, *p*_FDRc_=0.030. Interestingly, the LC displayed minimal covariance with most other regions; its BP_ND_^0^ appeared relatively independent, implying that noradrenergic-dopaminergic interactions in older adults may follow a distinct pattern from that of the more canonical nigrostriatal or mesolimbic pathways. Finally, SN and VTA were positively correlated with one another ρ(31)=0.481, *p*_uncorr_=0.005, *p*_FDRc_=0.039.

These cross-regional relationships in cross-sectional interindividual differences in receptor densities in ageing reinforce the notion that DA system integrity could follow anatomical or functional circuits (e.g., striatal loops, limbic circuits), rather than reflecting a uniform global measure. The overall pattern further highlights the nontransient trait-like nature of BP ^0^ in older adults, with distinct subcortical circuits, e.g. striatal versus midbrain versus limbic-thalamic, showing semi-independent profiles of D2/D3 receptor availability.

### Exploratory Factor Analyses of Interindividual Patterns in Baseline D2/D3 Receptor Availability

To explore whether the cross-ROI correlations reflect an underlying structure, principal-axis factoring (PAF) analysis with promax rotation was performed to the same averaged BP_ND_^0^ (detailed statistical diagnostics are described in the method section and Supplementary Method 5).

After extraction and rotation, the final two-factor solution explained 53.92% of the variance. Factor_1, accounting for 36.09%, showed the highest loadings in amygdala (0.469), hippocampus (0.562), LC (0.338), SN (0.777), thalamus (0.723), and VTA (0.740), echoing the components of the same limbic-midbrain-thalamic relationships found in the cross-ROI correlational analysis. Factor_2, accounting for 17.83%, was defined by the caudate (0.651), putamen (0.584), and NAcc (0.510), replicating the striatal cluster identified earlier by the significant caudate-putamen and putamen-NAcc correlations (Figure 3 and Supplementary Table 5). Intriguingly, SN showed a significant negative loading on the striatal factor. This result most likely suggests that this structure may have distinct functional affiliations itself, as the negative loading simply indicates how SN contrasts from striatal components in this dimension rather than a biologically inverse relationship.

**Figure 3.**
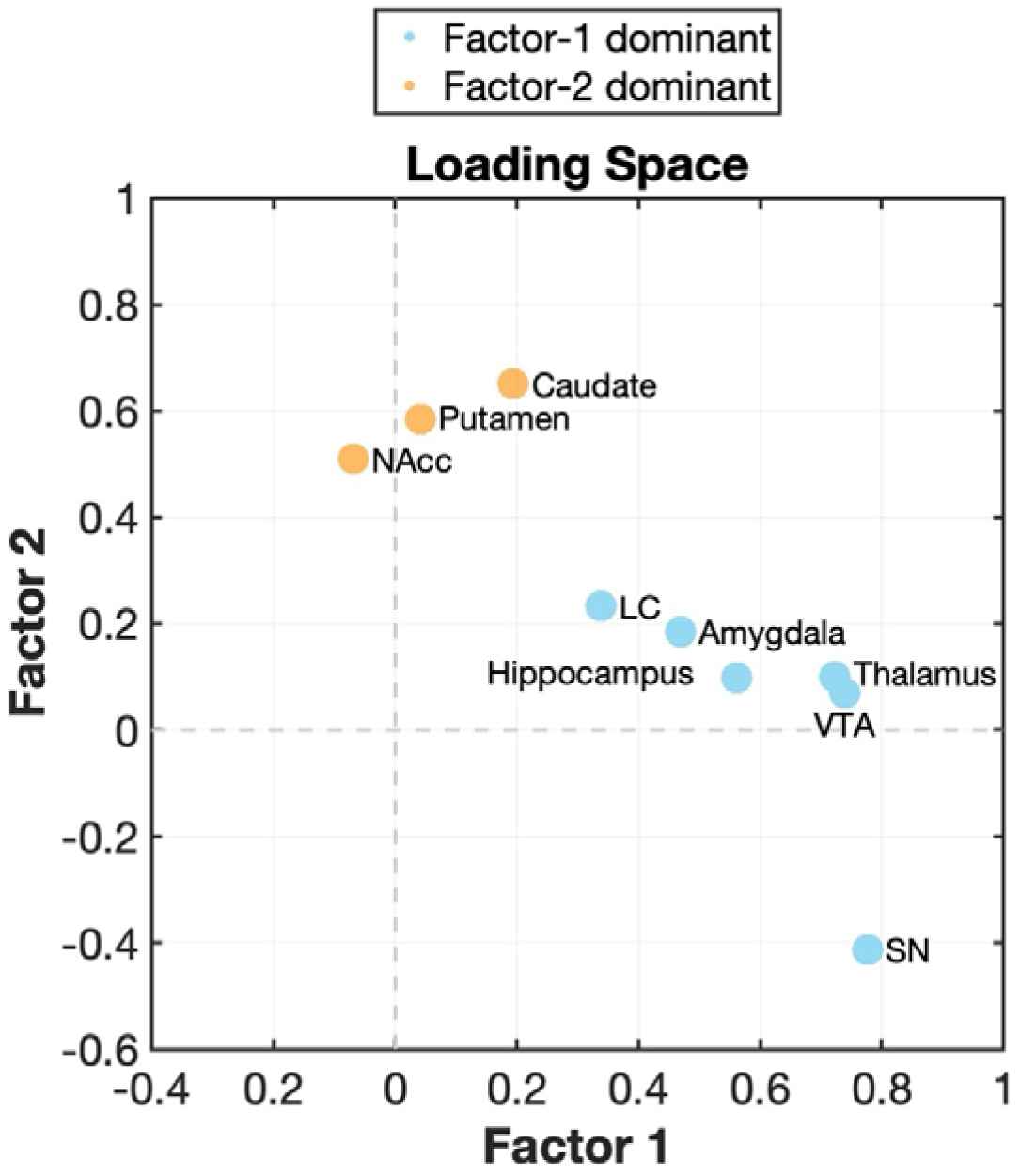
Loading plot from PAF (promax rotation) of session-averaged BP_ND_^0^ across nine dopaminergic ROIs. Each point represents a region’s pattern-matrix loading on Factor 1 (x-axis) and Factor 2 (y-axis). Blue dots denote ROIs whose highest loading was on the mesolimbic/mesohippocampal factor (Factor 1: LC, hippocampus, amygdala, thalamus, SN, VTA), whereas orange dots denote ROIs whose highest loading was on the striatal factor (Factor 2: caudate, putamen, NAcc). Dashed lines delineate zero loading; values farther from the origin indicate greater contribution to the corresponding factor. The plot illustrates the clear separation between the dorsal-/ventral-striatal cluster (upper left quadrant) and the extrastriatal mesolimbic cluster, with SN loading positively on Factor 1 and negatively on Factor 2.

**Figure 3.**
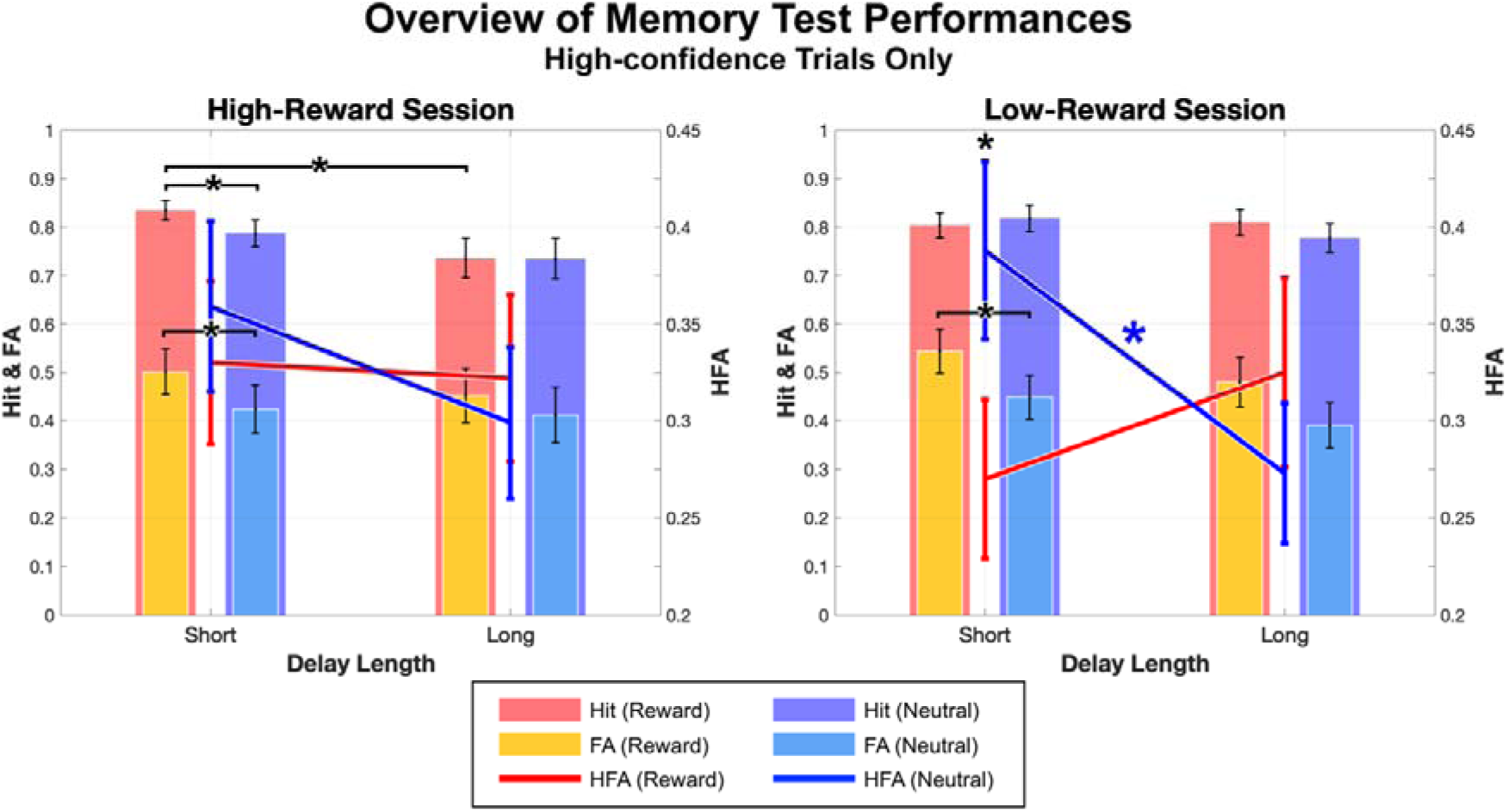
Overview of Recognition Memory test performance in trials rated with high confidence (‘sure’ compared to ‘not sure’). This figure compares the compound score of hit rate and FA (Hit - False Alarm), i.e. HFA (line graphs), as well as hit rate and FA (nested bar graphs) across two memory delay lengths, short and long, for reward-associated (warm colours) and neutral scenes (cool colours) within each reward context only for *high-confidence trials*. The left plot depicts high-reward sessions, and the right plot represents low-reward sessions, illustrating a distinct interaction between the reward context and delay length on overall memory performance. Whiskers around the marker in this plot represents standard error and an asterisk (*) represents statistical significance of *p*<.05. Again, the memory performance of low-reward sessions’ reward-associated scenes show an atypical pattern across temporal delays, and this pattern seems to be driven by the significant difference between FA for short-delay memory test in reward-associated and neutral scenes.

Together with the cross-ROI relationship patterns, this result supports a profile in which dopaminergic integrity in healthy older adults is organised around two semi-independent subsystems, i.e. a striatal factor reflecting dorsal and ventral striatum (caudate, putamen, and NAcc), and a mesohippocampal- or mesolimbic-factor encompassing ascending projections from midbrain neuromodulatory nuclei (SN, VTA, and LC) to limbic and thalamic targets.

### Relationship Between ROI Volume and D2/D3 Receptor Availability

As summarised in Supplementary Figure 5, unexpectedly, the nine volume and session-averaged BP ^0^ correlation pairs showed no significant correlations under either age-controlled or uncontrolled analyses.

No region showed a statistically significant correlation between volume and BP_ND_^0^ once multiple comparison correction was applied except for LC, which showed showed the largest zero-order correlation, but this association did not survive false-discovery adjustment (Supplementary Figure 5). This result suggests that, within our senior cohort sample, between-participant variation in D2/D3 receptor availability is not explained by a simple linear scaling with age-related macroscopic atrophy. The absence of a volume-BP relationship likely reflects the heterogeneous nature of brain ageing such as regional atrophy, microstructural change and receptor loss can proceed at different rates and be driven by partially independent mechanisms ^4,40^. Further discussion of these tentative patterns follows in the Discussion section below.

### Memory Performance Over Time and Across Reward Contexts

Analyses of recognition memory (Hit - False Alarm rate, denoted as **HFA** from here on) in trials rated with high confidence revealed a significant interaction between reward salience and delay length. Restricting analyses to high confidence responses isolates recognition based decisions and filters out low confidence noise, ensuring that effects reflect the strongest subjective memory traces^41^. Moreover, all key outcomes from the analysis including all trials remain virtually identical in magnitude and significance (Supplementary Result 2), confirming that the high confidence subset improves interpretability without altering core effect sizes. A three-factor repeated measures ANOVA (factors: Reward [Reward/Neutral], Delay Length [Short/Long], and Session Context [High-Reward/Low-Reward]) showed a significant Reward × Delay interaction, *F*(1,32)=9.435, *p*=0.004, η²=0.228. That is, initially in short-delay tests, reward-associated scenes showed worse memory discriminability than neutral scene (Figure B). However, after the 24-hour delay, performance for reward and neutral scenes converged, suggesting that any immediate disadvantage conferred by reward salience in the short term dissipated over time. Contrary to our hypothesis, no significant main effect of reward was observed, *F*(1,32)=0.474, *p*=0.496.

A follow-up analyses of **hit rates** in high-confidence trials similarly showed the influence of delay length and reward context. There was a main effect of Delay Length, *F*(1,32)=6.824, *p*=0.014, with significantly higher hits in short-delay tests compared to long-delay tests, as expected. In addition, there was a trend indicating that the low-reward sessions showed higher hit rates than high-reward sessions, *F*(1,32)=3.001, *p*=0.093. A three-way interaction showed trend-level significance, *F*(1,32)=3.795, *p*=0.060, η²=0.106, suggesting that, under high-reward contexts, reward-associated scenes had higher hit rates than neutral during short-delay tests, but this advantage did not persist after the 24-hour delay. Moreover, there was a further trend (*<u>F</u>*[1,32]=3.929, *p*=0.056) indicating that reward-associated scenes at long delay were better remembered in low-reward sessions than in high-reward sessions, potentially reflecting a distinct influence of reward salience at encoding stage depending on the motivational context, albeit with an advantage for the low-reward context.

In another follow-up three-factor repeated measures ANOVA, false alarms (**FA**) in trials rated with high-confidence were significantly influenced by reward salience, *F*(1,32)=7.478, *p*=0.010, with reward-associated scenes showing significantly higher FA than neutral scenes. This pattern was most marked in the low-reward contexts, where reward-associated scenes showed higher FA compared to neutral scenes, *F*(1,32)=5.457, *p*=0.026, whereas high-reward sessions showed a smaller and statistically nonsignificant difference between the two types of scenes. This result suggests that the unexpected absence of the anticipated reward effect on memory performance, HFA, primarily arises from a reward-adverse effect on FA.

Finally, when high- and low-reward context sessions were averaged across, a two-factor repeated measures ANOVA (factors: Reward and Delay Length) in high-confidence trials revealed a main effect of Reward at a trend-level significance, *F*(1,32)=3.578, *p*=0.068, with neutral scenes showing higher HFA, at short delay in particular. On the other hand, reward-associated scenes were again associated with significantly higher FA, *F*(1,32)=6.804, *p*=0.014.

Overall, these results show (1) that memory discriminability declines over time mainly due to declining hit rates; (2) that reward-associated scenes are subject to increased FA; and (3) that the impact of reward salience may depend critically on motivational context.

### Relationship Between Baseline Receptor Availability and Memory Performance

Nonparametric partial correlation analyses using Spearman’s rank-order examined how individual differences in D2/D3 receptor availability related to different aspects of memory performance in trials with stronger subjective memory trace (high-confidence HFA, hit rates, and FA), while controlling for age. Given the known and expected effect of reward context on behavioural performance^42^, in this analysis, we treated each experimental session as an individual datapoint (Figure 4; red dots indicating data from high-reward session and blue ones from low-reward session). This approach was chosen because both the memory test scores and BP_ND_^0^ values were obtained separately for each session. Although no statistically significant overall session effect was observed for BP_ND_^0^ (Figure 1), analysing session-level data preserved the granularity of the measurements and allowed for a more precise alignment between memory performance and physiological state within each session. Given that several memory test performance indices showed session-specific patterns (Figure 3), the correlation between memory and BP_ND_^0^ was also examined separately for each reward context (high- and low-reward contexts) as a post-hoc analysis and to probe potential latent patterns in the relationship between DA integrity and memory.

**Figure 4.**
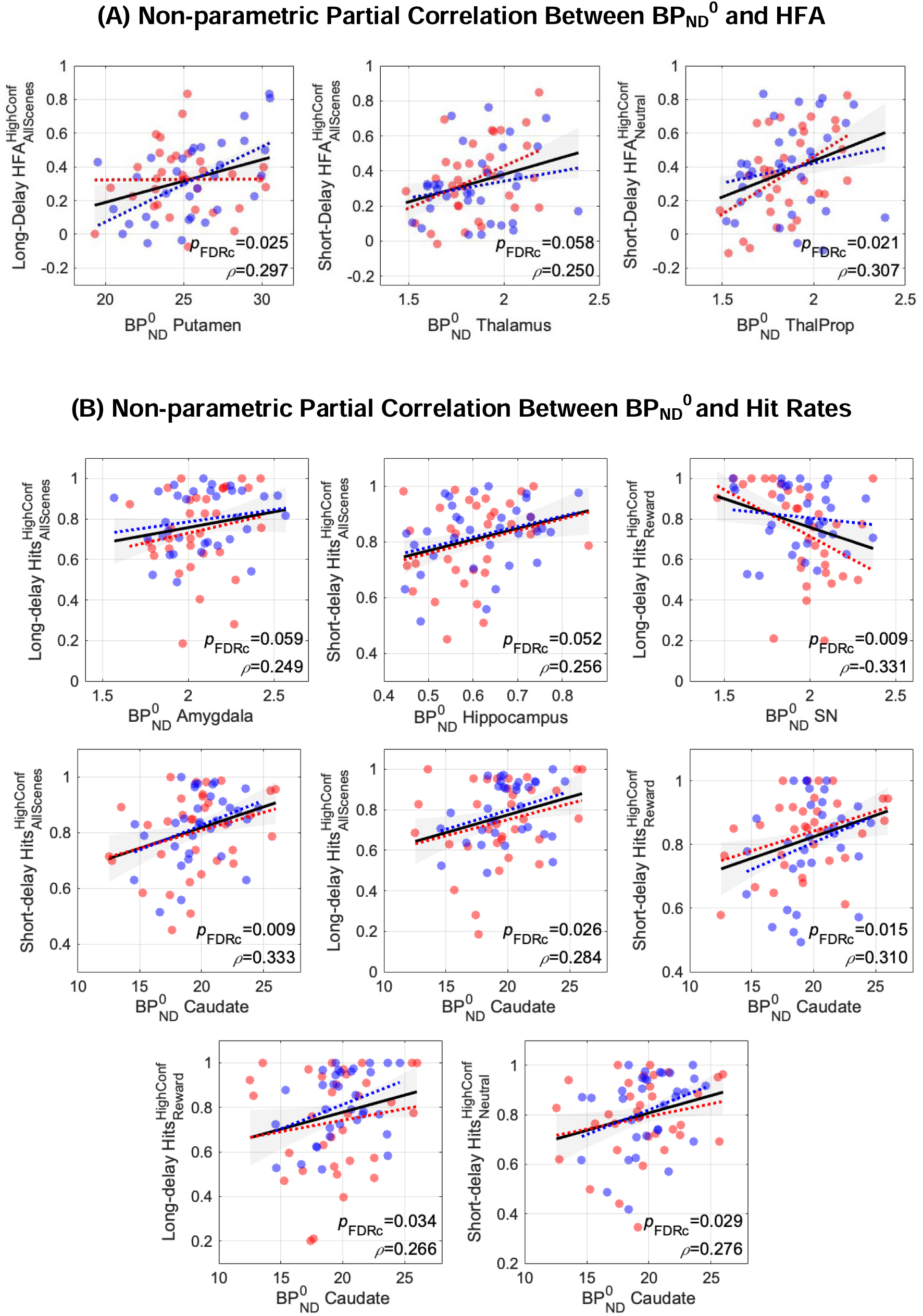
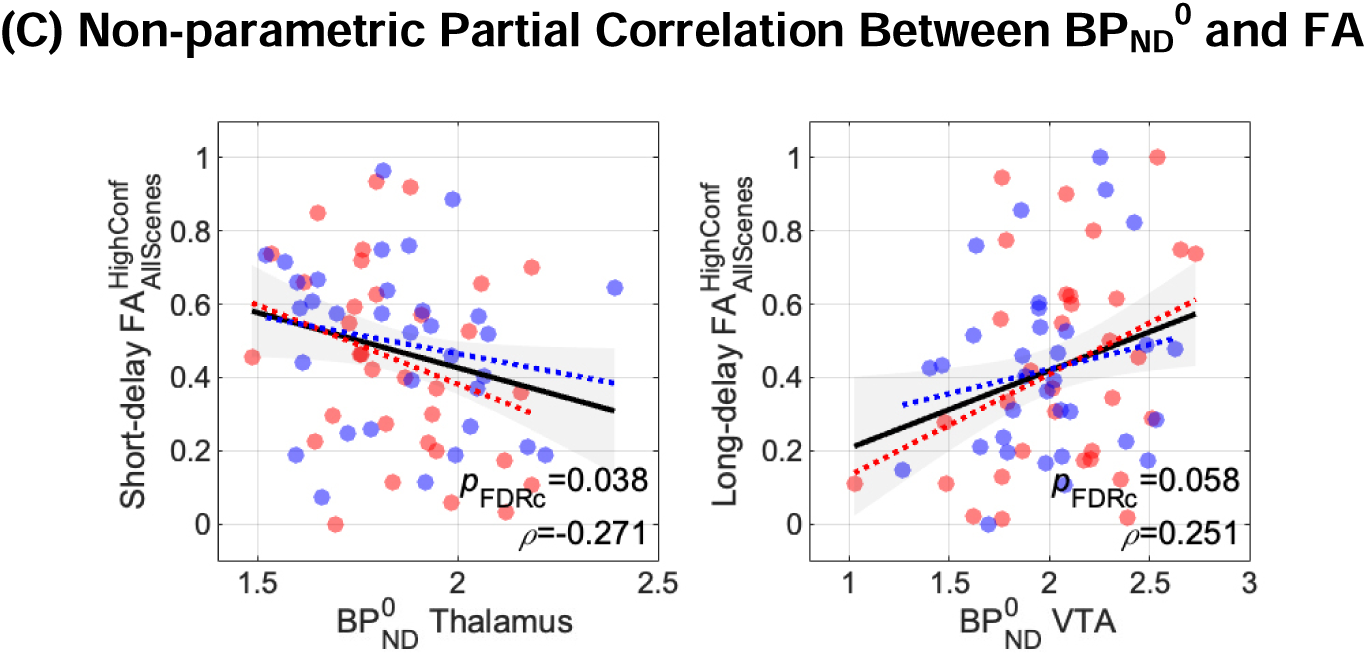
Age-controlled non-parametric correlation between baseline D2/D3 receptor availability and memory test performance indices. The above scatterplot figures illustrate the results of correlational analyses between BP_ND_^0^ measured in each ROI and memory test performance incides, i.e. HFA, hit rates, and FA, of subsequent incidental memory tests assessed at two delay intervals, short- and long-delay (∼15 minutes and 24 hours after encoding, respectively). Each dot represents a single data point, red dots indicating data from high-reward session and blue ones from low-reward session. Linear fits are drawn as a solid black line for both sessions’ data, a red dotted line for high-reward session’s data, and blue dotted line for low-reward session’s data. Grey shaded areas around the solid black line denote the standard error. Within each plot, the Spearman’s rho (ρ) and FDR-corrected p-value (*p*_FDRc_) are displayed in the bottom right corner. Analyses were conducted separately for: **(A) HFA**, (**B) hit rates**, and **(C) FA**.

Firstly, higher BP_ND_^0^ in the putamen correlated with superior long-delay discriminability (that is, Hit - False Alarm [HFA]) for all scenes, ρ(64)=0.297, *p*_uncorr_=0.016, *p*_FDRc_=0.025. In the thalamus, BP_ND_^0^ showed positive trend-level correlation with short-delay discriminability for all scenes, ρ(64)=0.250, *p*_uncorr_=0.044, *p*_FDRc_=0.058, as well as for neutral scenes, ρ(64)=0.307, *p*_uncorr_=0.013, *p*_FDRc_=0.021.

When the correlation between BP_ND_^0^ and high-confidence hit rate was examined, we found that subject who showed greater BP ^0^ in the caudate nucleus tend to show higher hit rates across most stimulus categories at both short delay and long delay. Significant BP_ND_^0^ correlations include positive correlation with short-delay hit rates for all scenes, ρ(64)=0.333, *p*_uncorr_=0.007, *p*_FDRc_=0.009, reward-associated scenes, ρ(64)=0.310, *p*_uncorr_=0.012, *p*_FDRc_=0.015, and neutral scenes, ρ(64)=0.276, *p*_uncorr_=0.026, *p*_FDRc_=0.029. In addition, greater BP_ND_^0^ was associated with long delay hit rates for all scenes combined, ρ(64)=0.284, *p*_uncorr_=0.022, *p*_FDRc_=0.026, and reward-associated scenes, ρ(64)=0.266, *p*_uncorr_=0.032, *p*_FDRc_=0.034. The hippocampus showed a trend-level positive correlation with short-delay hit rates, ρ(64)=0.256, *p*_uncorr_=0.040, *p*_FDRc_=0.052. BP_ND_^0^ in the amygdala showed positive correlation trend with long-delay hit rates, ρ(64)=0.249, *p*_uncorr_=0.045, *p*_FDRc_=0.059.

Unexpectedly, individuals with higher SN BP_ND_^0^ showed worse long-delay hit rates for reward scenes, ρ(64)=-0.331, *p*_uncorr_=0.007, *p*_FDRc_=0.009. Lastly, higher BP_ND_^0^ in the thalamus negatively correlated with short-delay FA, ρ(64)=-0.271, *p*_uncorr_=0.029, *p*_FDRc_=0.038, whereas, intriguingly, greater BP_ND_^0^ in the VTA showed a trend-level association with higher long-delay FA, ρ(64)=0.251, *p*_uncorr_=0.044, *p*_FDRc_=0.058.

## Discussion

### Stability and Regional Architecture of Baseline D2/D3 Receptor Binding in Ageing

Our findings demonstrate that baseline D2/D3 receptor availability (BP ^0^) in healthy older adults is in general a highly stable individual characteristic across sessions (Figure 1). There was no systematic difference in D2/D3 receptor availability between the high- and low-reward context sessions conducted weeks apart, indicating that anticipated reward had no immediate effect on resting D2/D3 receptor binding. The observed test-retest stability is in line with existing test-retest studies in older adults reporting that [^18^F]fallypride binding is reproducible across sessions, with high intra-class correlations (≥0.80) and low variability (<8%) in striatal and limbic regions^33^. In other words, inter-individual differences in dopaminergic receptor availability reflect relatively stable biological traits rather than transient state-to-state fluctuations. Such stability corroborates the notion that baseline D2/D3 receptor availability measures can serve as reliable biomarkers in ageing research.

We also observed, however, that test-retest reliability was slightly lower in certain regions, namely NAcc and putamen. These findings likely reflect region-specific factors rather than true receptor change. The putamen was shown to exhibit a steep rostrocaudal gradient of D2/D3 receptor density (∼20-25% higher binding in dorsal versus ventral putamen)^38^. Averaging across this axial gradient could compress between-subject variability in binding, thereby contributing to masking true individual differences^39^. Furthermore, as spontaneous movements such as bathroom trips after the In-flow scan, postural adjustments, or finger movements, are inevitable across our 6-hour experimental visit and vary idiosyncratically between days. As the putamen has dense DA terminals^43^ and therefore is more susceptible for short-term endogenous DA fluctuations that can be induced by even very simple 2-minute foot flexing movements and be captured by PET imaging, where such a simple motor task reduced [^11^C]raclopride BP in the putamen by up to 30%^37^. This element may have curtailed the interscan correlation in this region. Meanwhile, the NAcc is a relatively small region bordered by high-uptake areas (caudate, putamen) and low-uptake areas (cortex), making its PET signal prone to partial-volume spillover. This could undermine reliability and has been noted in previous PET studies where test-retest intraclass correlations for NAcc binding were often ≤0.60, compared to ≥0.80 in larger striatal regions^33,35,36^. In summary, these factors might explain the minor deviations from otherwise strong cross-session stability, rather than indicating any rapid biological change in receptor densities. However, further research is needed confirming these assumptions.

In addition to being stable within-subject, baseline D2/D3 receptor binding exhibited a complex regional architecture across the brain. Our exploratory factor analysis using PAF implies that dopaminergic integrity in ageing may not be uniform: we found an emergent separation between a striatal factor (dorsal and ventral striatum: caudate, putamen, NAcc) and an mesolimbic factor (limbic, thalamic, and midbrain regions). That is, an older adult with relatively high D2/D3 receptor availability in striatal regions did not necessarily have high D2/D3 receptor availability in midbrain or limbic regions. This partial dissociation is consistent with evidence that age-related DA declines are regionally heterogeneous^40^. For example, cortical and hippocampal D2/D3 receptors can exhibit different ageing trajectories than striatal receptors, and dopaminergic losses do not strictly mirror the spatial pattern of gray matter atrophy or global cognitive decline^40^. Our findings from PAF extend this idea by suggesting that dopaminergic ageing in certain pathways (e.g. striatal vs. mesolimbic or thalamic circuits) may occur somewhat independently. Striatal D2/D3 receptor integrity might be relatively preserved in some individuals who nevertheless show lower limbic or midbrain receptor binding, or vice versa. These regional covariance profiles highlight the importance of examining pathway-specific DA changes instead of treating the older adult’s dopaminergic system as a monolithic entity.

We also found that individual differences in regional brain volume were largely unrelated to D2/D3 receptor binding in those same regions. None of the nine ROI volumes showed a significant correlation with their D2/D3 receptor availability after correcting for multiple comparisons (see Supplementary Figure 5). This converges with prior observations that the topography of DA receptor loss with age does not scale in direct proportion to regional atrophy. Some areas show pronounced receptor decline despite limited volume loss, whereas others show the opposite pattern^40^. Dopaminergic integrity in ageing thus appears to carry unique variance reflecting neurochemical decline beyond what is captured by morphology observed via MRI. This dissociation is encouraging for PET studies because it suggests PET-measured D2/D3 receptor changes are not merely epiphenomena of brain shrinkage but rather represent a distinct aspect of brain health. Viewed within current theories of cognitive ageing, our findings are consistent with the brain maintenance hypothesis, the idea that some older adults manage to maintain levels of neurochemical function comparable to those of young adults despite age-associated changes in the brain^44^. For example, a person may preserve a high density of DA receptors (supporting cognition) even if they exhibit some cortical thinning or volumetric loss. The strong stability and the regional specificity of D2/D3 receptor availability observed in our study accentuate the need to consider multiple dopaminergic subsystems when linking DA to behaviour in older adults. We note, however, that the present cross-sectional dataset cannot distinguish genuine maintenance from trait differences established earlier in adulthood nor does it rule out more complex or age-dependent coupling between structural loss and receptor decline. Longitudinal or lifespan datasets will be required to determine whether such links emerge earlier in adulthood or during periods of accelerated ageing.

### Reward Salience and Episodic Memory in Older Adults

In addition to imaging, our study examined how motivational context (high- and low-monetary reward contexts) influenced episodic memory in ageing. Behaviourally, the effects of reward on memory were mixed and depended on the experimental context and retention delay. We found that high-reward contexts did not uniformly boost memory for all participants or across all test delays. Instead, benefits of reward on recognition memory appeared selectively. For instance, in high-reward sessions, there was minimal difference in discriminability performance (HFA) between the reward conditions as well as across two retention delays. However, in low-reward contexts, reward-associated scenes were actually remembered worse than neutral scenes on the short-delay tests (i.e. 15 minutes after the end of encoding phase), whereas, after a 24-hour delay, marginal performance increase for reward-associated scenes emerged while the performance for neutral scene significantly decreasing, converging the performance level of neutral and reward-associated scenes comparably (Figure 3, left panel). These findings suggest that the mnemonic advantage of reward salience in older adults is conditional, and the conditions under which a reward benefit or ‘cost’ appears may involve how salient the overall motivational context is and whether memory is tested immediately or after a consolidation period.These complex effects align with prior research showing that motivation can improve memory, but that ageing alters this relationship. Some studies report that older adults, like young adults, gain a memory advantage for high-value or rewarded items^45^. In those cases, anticipation of reward improves encoding, suggesting that the fundamental dopaminergic mechanisms of reward motivation remain intact in healthy ageing^18,45^.

On the other hand, other work finds that older adults do not benefit as much as younger adults from reward incentives, especially for long-term episodic retention^46^. For example, Geddes et al. (2018) observed that younger adults had significantly better 24-hour recall for reward-paired items (specifically via enhanced recollection), whereas older adults showed no such memory gain. Our results align with this literature. The presence of a reward effect in some conditions suggests that older adults can still engage motivational pathways to bolster memory when task conditions favour it (e.g. when given time to consolidate or when reward cues are salient). At the same time, the absence of a broad, uniform advantage of reward in our results, particularly in short-delay memory tests, may reflect age-associated decline in how effectively neuromodulatory reward signals are integrated during encoding. In particular, the Reward × Delay Length interaction in our data is consistent with theories that DA-driven benefits primarily arise for long-term memory and are less integral for short delays^47,48^. Indeed, older adults in our study did not show a robust overnight improvement for reward items, but the elimination of the initial deficit by 24 hours suggests that the detrimental effects of reward on immediate memory did not carry over into long-term retention and dopaminergic engagement during learning is thought to support memory persistence specifically beyond a few hours^49^.

Thus, the pattern observed in our older sample, namely reward-related memory after longer delays compared to shorter ones, suggests that reward-related consolidation processes, which is likely DA-mediated, remain functional, even if immediate encoding advantages are far subtler. As this delay-specific pattern was not pre-hypothesised, we regard it as exploratory and in need of replication. In summary, we interpret the overall behavioural findings as evidence that reward motivation can aid episodic memory in ageing but its efficacy is conditional, possibly requiring optimal engagement of DA-dependent consolidation processes or particular task settings to observe more conspicuous benefits.

### Linking DA System Integrity to Reward and Memory

The pattern of correlations between baseline D2/D3 receptor availability (BP_ND_^0^) and recognition performance highlights a complex influence of DA in ageing. Across participants, a larger receptor pool in any single ROI did not translate into a uniform mnemonic advantage. Instead, the direction and magnitude of associations differed with across ROIs and behavioural indices (i.e. hit rates, FAs, or the compound HFA measure). This dissociation can be informative. High hit rates indicate efficient detection of studied items or may imply liberal response tendency due to stronger subjective memory trace, high FA mark a liberal endorsement of lures, and HFA gauges the fidelity with which old and new scenes are distinguished. The coexistence of positive and negative associations across regions therefore suggests that DA can facilitate certain elements of recognition memory while constraining others, depending on the circuit that is engaged and the mnemonic component that is probed.

In our correlational analyses, positive associations were observed in several other dopaminergic ROIs with memory test performance and point to DA’s facilitatory potential in healthy ageing. High D2/D3 receptor availability in the thalamus, for instance, predicted fewer short-delay HFA, especially for neutral scenes, highlighting that better immediate discriminability might depend on dopaminergic integrity in the thalamus, a region involved in attentional gating and sensory relay^50,51^. Thus, older adults with a well-preserved thalamic dopaminergic system might be more efficient at gating and transmitting information during encoding, leading to fewer FA for non-salient material. Intriguingly, the advantage of high receptor availability in the thalamus was restricted to the short-delay tests only. This pattern suggests that the benefit may stem from encoding- or early-retrieval-related processes rather than a static dopaminergic tone, because overnight consolidation likely engages additional, wider networks. At the same time, higher D2/D3 receptor availability in the thalamus predicted fewer FA for all scene types during short-delay tests (Figure 4C), suggesting that older adults with preserved dopaminergic integrity in the thalamus may more effectively encode and reject unstudied scenes. Again, this finding is consistent with the thalamus’ established role as a sensory relay and attentional gate in memory encoding^50,51^.

Moreover, the dorsal striatum (caudate and putamen) yielded a robust pattern of positive correlations with hit rates at both short and long delays, particularly for reward-associated scenes in the low-reward session (Figure 4A, 4B, and Supplementary Figure 7A-2 and 7B-2). This finding indicates that, even when external incentives are modest, dopaminergic integrity in the dorsal striatum can bolster performance during recognition tests, possibly by increasing executive control of retrieval searches or by adjusting decision thresholds that govern ‘old vs. new’ responses^44,57,58^. However, considering the overall pattern, this finding adds more nuance to the interpretation. Specifically, D2/D3 receptor availability in the caudate did not coincide with higher HFA (cf. Figure 4A and 4B) but only with higher hit rates and, in session-specific analyses, correlated positively with long-delay FA when motivational context was high (Supplementary Figure 7C-1). The most parsimonious interpretation of these pattern of findings could be that a richer dopaminergic environment in the caudate facilitates a more liberal response tendency, raising both correct recognitions and lure acceptances, rather than genuinely enhancing the memory. In contrast, subjects who showed higher D2/D3 receptor availability in the putamen also tend to show higher long-delay HFA across both sessions (Figure 4A; and also with reward-scene HFA in low motivational context [Supplementary Figure 7A-2]), implying genuine enhancement of memory is accompanied by low FA. The putamen’s dense sensorimotor projections and role in Pavlovian control^59^ as well as in anticipated actions^60^ make it a reasonable substrate for stabilising encoded information over longer temporal delay^61^. Taken together, these results suggest a functional dissociation within the dorsal striatum regarding memory processes and behaviour. Dopaminergic integrity in the caudate appears to modulate response policies, whereas that in the putamen supports the durability of memories.

In a related vein, a separate session-specific analysis, subjects with higher D2/D3 receptor availability in the LC of low-reward session showed better short-delay hit rates for reward-associated scenes (see Supplementary Results 7B-2). This effect was not found in the cross-session analysis, for HFA rates, or for long-delay memory test performance, suggesting that a more responsive LC may help older adults steer behaviour toward motivationally relevant information when overall motivational salience is modest.

Although the correlations in the amygdala, hippocampus and VTA did not meet the corrected threshold, their directions map closely onto established circuit functions and thus merit consideration. D2/D3 receptor availability in the amygdala showed a positive trend with long-delay hit rates for reward-associated scenes (Figure 4B). Animal and human work indicates that dopaminergic activation of basolateral amygdala during or shortly after learning enhances late hippocampal plasticity and stabilises emotionally or motivationally salient memories ^42,47^. A more robust dopaminergic integrity in this region in older adults may therefore preserve this modulatory pathway, yielding moderately better 24-hour retention. Furthermore, the hippocampus showed a trend-level positive correlation with short-delay hit rates (Figure 4B). This observation is consistent with the hippocampus’s central role in forming and retrieving detailed episodic representations, a process that is modulated by DA^20^. Human fMRI studies report hippocampal activation during 20-second object-location binding tasks^62^ and during delayed matching of novel scene layouts^63,64^, demonstrating that hippocampal integrity is required for associative memory well under the 24_h timescale. Although the effect in our cohort did not survive multiple-comparison correction, its direction aligns with these mechanistic accounts and suggests that preserved hippocampal D2/D3 receptors may support the fidelity of early episodic encoding in older adults.

In the VTA, on the other hand, higher D2/D3 receptor availability showed a modest tendency with more long-delay FA without any hit-rate relationship (Figure 4C). The observation that higher D2/D3 receptor availability in the VTA predicts more long-delay FA, without any accompanying rise in hit rates or HFA, argues against a simple liberal bias as we observed in the caudate-hit rate relationship above. Instead, it points to a mnemonic error, an increased tendency to misattribute familiarity to novel lures. The VTA is known to be strongly engaged by novel stimuli and conveys a salience signal to its projection areas^65,66^. If individuals with higher tonic VTA receptor availability generate stronger novelty signals upon reencountering lures, they may experience those items as spuriously familiar and thus endorse them. In ageing, this mechanism may be exaggerated, so that individuals with higher tonic VTA receptor availability show impaired mnemonic discrimination at long delay, manifesting as isolated FA. In this light, the VTA-FA association might reflect a dopaminergic shift toward familiarity-driven retrieval rather than a change in overall response criterion.

Finally, intriguingly, a strong negative correlation was found for the SN. Older adults who showed high D2/D3 receptor availability in the SN also showed the poor hit rates for reward-associated scenes after the 24-hour delay. One possible mechanistic explanation is that abundant D2/D3 autoreceptors^71^ in this region may restrict the phasic DA bursts, thereby attenuating the consolidation boost that follows reward delivery^16,20^. Stable, characteristical BP in midbrain D2R density has been documented with PET, and individuals at the upper extreme show a more rigid dopaminergic response profile^72^. In line with this, Richter et al. (2017) demonstrated that individuals genetically predisposed to lower D2 receptor expression outperformed high-expressing counterparts in 24-hour recognition of reward stimuli, suggesting that D2 *excess* can actually impair memory for stimuli with high motivational value^73^. This evidence ties with our results linking higher D2/D3 availability in the SN to worse long-delay recognition test performance for reward-associated scenes and further supports the notion that there is an optimal, rather than a maximal, dopaminergic tone for the consolidation of motivationally relevant information^74^. This finding supports an inverted-U hypothesis in which excessive dopaminergic neuromodulation, excitatory or inhibitory, can hinder the reinforcement of salient traces^49,74–76^. Although additional experiments are needed to confirm this explanation, several findings in this study suggest that dopaminergic health in older adults is not a simple ‘the more, the better’ phenomenon, and that certain individuals with high SN D2/D3 receptor availability might be particularly vulnerable to suboptimal memory outcomes for stimuli with higher motivational salience.

Our volume-BP correlations add another layer to this complex landscape. No relationships between ROI volumes and their respective D2/D3 receptor availability measure (BP ^0^) survived multiple-comparison correction. This suggests that morphometric preservation (represented by TIV-normalised volume) and dopaminergic integrity (represnted by BP_ND_^0^) may diverge substantially in the process of healthy ageing. Because no strong volumetric-BP connections emerged, the correlations with memory cannot simply be attributed to structural deterioration. It is plausible that dopaminergic receptor availability, being relatively stable over the sessions and less conflated by partial volume effects, provides an independent index of neuromodulatory system health that impacts recognition accuracy.

Importantly, the stability of BP ^0^ measure across the high- and low-reward sessions suggests to that these DA-memory correlations likely capture relatively stable differences in the dopaminergic system rather than transient states. Older adults in our sample showed reliable individual cross-session in D2/D3 receptor availability, reinforcing the view that dopaminergic health is a lasting characteristic rooted in each person’s neurobiology^33^. Therefore, whether DA confers greater short-delay recollection or more frequent FA to salient scenes, it appears to do so consistently for a given individual over time. However, since memory performance varied more across sessions, these associations should be interpreted with caution and viewed as potentially context dependent. This stable property is methodologically advantageous, as it reduces the risk that session-specific fluctuations might obscure real underlying relationships with memory processes. It also implies a target for intervention studies. If those with the highest SN or VTA D2/D3 receptor availability consistently show less robust memory performance for items associated with reward salience, it might be worthwhile to test DA-stabilising approaches in that subgroup to examine if memory outcomes can be improved in a longitudial or pharmacological follow-up design.

## Study limitations

This study is not without caveats. First, despite carefully planning the timing of the In-Flow, Baseline, and Task scans, the baseline phase did not permit full target-to-reference equilibrium in the highest-binding striatal ROIs (caudate, putamen, NAcc). Consequently, BP ^0^ values in these regions may be underestimated and should be interpreted cautiously relative to extrastriatal values. However, simulations and empirical work with [¹_F]fallypride show that reducing an 180-minute [¹_F]fallypride scan *step-wise* by up to 60 minutes lowered BP_ND_ estimated with SRTM in the putamen by only ≈1% per ten-minute block, and, crucially, left between-subject rank order virtually unchanged^32^. Hence absolute values may be biased downward, but the individual-difference signal we correlate with behaviour is largely preserved.

Secondly, our sample of 33 older adults is modest and the inter-session interval varied (*M*±*SD*≈6±9 weeks), which limits power to detect subtle effects and could introduce session-specific noise. Even so, this cohort size is typical for dual MR-PET ageing studies, and our two-session design yielded 60 usable session-level datapoints after accounting for drop-outs. Importantly, a dedicated test-retest investigation that scanned older adults with a three-block [¹_F]fallypride protocol 4 to 6 weeks apart reported intraclass correlation coefficients ≥0.80 in both striatal and limbic ROIs^33^, indicating that receptor rankings remain highly stable over intervals comparable to ours.

Thirdly, all analyses are correlational; causality between baseline dopaminergic system health and memory cannot be inferred. Nonetheless, while causality cannot be claimed, the majority of the same ROI-memory associations replicate across (1) two motivational contexts and (2) short-vs long-delay tests, lending convergent validity.

Fourth, multiple comparisons meant many ROI-behaviour associations did not survive FDR correction, raising the possibility of Type I errors. Indeed, FDR correction pruned many effects, yet the surviving findings align with anatomically plausible circuits (striato-thalamic and LC networks) that echo independent PET and pharmacological findings, reducing the probability of Type-I error.

Finally, the monetary incentive was deterministic and symbolic (€120 vs €15, each reward-associated trials being rewarded at 100% chance) rather than performance-contingent, which could have muted reward salience and thereby phasic DA responses for some participants. However, our choice of deterministic (100%) reward probabilities was intentional in order to eliminate variance from cognitive processes such as risk evaluation and expectancy violation^77,78^, each with its own neurochemical footprint (NA, acetylcholine). This choice allows us to relate baseline D2/D3 receptor availabiity to pure expected-value signals without confounds from surprise- or risk-related neuromodulation.

Together, these caveats temper, but do not overturn, our conclusions and point to concrete improvements for future studies of motivated memory in ageing. Future work with larger samples, longer acquisition protocols, and more robust experimental manipulations and cognitive task could address these challenges.

## Conclusion

In conclusion, our findings indicate that higher D2/D3 receptor availability in older adults is not invariably beneficial for recognition memory, and that reward contexts do not uniformly modulate these relationships. Instead, each measure of memory, hit rates, FA, and HFA, reveals a unique perspective on how the integrity of the dopaminergic system intersects with both salience and retrieval intervals in ageing. Behaviourally, reward’s impact on recognition varied with retention interval and context, with reward-associated scenes ‘recovering’ from an initial disadvantage suffered in short-delay recognition tests by 24 hours. Moreover, from the examination of relationship between memory and D2/D3 receptor availability, SN and VTA appear to correlate negatively with reward-associated long-delay memory performance, amygdala, putamen, hippocampus, and thalamus positively with both short- and long-delay memory indices, whereas the caudate appeared to contribute to more liberal response bias. A factor analysis revealed two partially independent systems, one centred on the striatum (caudate, putamen, NAcc) and a second comprised of mesohippocampal/mesolimbic regions, suggesting that dopaminergic integrity may decline in a regionally specific manner. No explicit relationship emerged between regional volumes and D2/D3 receptor availability, showing that receptor density provides neurochemical insight beyond volumetric deterioration in ageing measured by structural MRI. Together, these findings show that the ageing DA system contributes to memory in a heterogeneous, circuit-specific manner, shapes an individual’s ability to leverage reward salience in mnemonic processes, bearing implications for more targeted interventions in memory ageing.

## Methods

### Participants

Thirty-three healthy older adults (13 female; age: 64-85 years, *M*±*SD*=71.48±5.52) were recruited through the DZNE memory clinic database and local advertisements. Exclusion criteria included psychiatric or neurological disorders, psychoactive medication use, or contraindications for PET. All participants demonstrated normal cognition (MMSE^79^>=26) and minimal depressive symptoms (BDI^80^<15). Written informed consent was obtained >=48 hours prior to tracer application. The study was approved by the local Ethics Committee and the Federal Office for Radiation Protection (BfS).

### Experimental Design and Procedure

Each participant underwent two MR-PET sessions, separated by ∼42 days (*M*±*SD*=42.2±63.3), each corresponding to a high- or low-reward condition. Before scanning, participants completed practice tasks to familiarise themselves with the button-press responses and reward contingencies. During the MR-PET session, participants received an intravenous bolus of [^18^F]fallypride and, after an initial baseline phase, performed a ∼50-minute reward-based categorisation task inside the scanner. The amount of monetary reward differed between the two sessions (high vs. low), with reward-associated categories predefined (indoor vs. outdoor images; public vs. private for indoor set; urban vs. natural scenes for outdoor set). Immediately following scanning, participants completed a short-delay (∼15-minute) recognition memory test. A long-delay (24-hour) memory test was administered on a subsequent visit, with separate sets of old and new images. All visual stimuli were drawn from a controlled pool^81^, screened for memorability, emotional content, faces, readable text, and animals. Luminance was controlled to 50% in all images. Further details on experimental protocols are provided in the Supplementary Method 1.

### MR-PET Acquisition and Preprocessing

All imaging was conducted on a 3T Siemens Biograph mMR scanner with a 24-channel head coil. PET data were acquired over ∼175 minutes, starting with a 60-minute in-flow dynamic scan (60×1-minute frames), followed by a 15-minute baseline scan and a 55-minute task scan. [^18^F]fallypride synthesis followed established protocols^82^, and quality control ensured >97% radiochemical purity and specific activities >8 GBq/mmol. More detailed procedure for PET reconstruction protocols is described in the Supplementary Method 2. Two T1-weighted (T1w) MPRAGE images (1 mm isotropic) were acquired per session for anatomical reference, and double-echo fieldmaps were obtained for EPI distortion correction. Functional MRI was collected with a T2*-weighted, gradient-echo EPI sequence (2mm isotropic voxels, TR=3600ms, TE=32ms). A full list of structural sequences (e.g., T2, FLAIR, MTC) is in the Supplementary Method 2, as these measures do not form the primary focus of the present analyses.

Two T1w images per subject were bias-field corrected and segmented into grey matter, white matter, and CSF using SPM12’s Segment function. To improve segmentation near the cerebellar cortex, the Bias FWHM parameter was adjusted to 30mm. The corrected T1w images were then processed with Freesurfer’s recon-all (v7.4.1, with the - all flag) for whole-brain segmentation, which was subsequently rigidly registered back to native T1w space and was visually inspected for segmentation errors specifically in the cerebellar cortex, this study’s reference region to minimise errors in pharmacokinetic modelling. Subcortical segmentations of SN and VTA were extracted from the probabilistic atlas by Pauli et al. (2018) using FSL’s fslmaths, while LC segmentation followed the biologically referened mask from Dahl et al. (2022). Each was non-linearly transformed to native T1w space via antsRegistrationSyN.sh and antsApplyTransforms (nearest-neighbour interpolation), using a study-specific structural template as an intermediate. Finally, the SN, VTA, and LC segmentations were integrated with native-space whole-brain segmentations using a custom MATLAB script. Further transformation details and exhausted steps for selecting and delineating ROIs are provided in Supplementary Method 3 and Supplementary Table 1.

### PET Data Preprocessing and Pharmacokinetic Modelling

PET frames were rigidly registered to native T1w space, and time-activity curves extracted from ROIs (for the details of the ROI selection and delineation process, please refer to the previous item and Supplementary Method 3). Initially, ROIs included DA source areas (SN, VTA, LC) and target areas (thalamus, amygdala, hippocampus, and striatal regions), selected based on D2/D3 receptor density and relevance to ageing and memory ^8,83^ (Supplementary Method 3). Although putamen, caudate, and NAcc did not reach kinetic equilibrium in our scans^84,85^ (Supplementary Figure 3 and Supplementary Table 2), their Baseline BP_ND_ (BP_ND_^0^) can still offer insight into overall receptor availability (more detailed rationale is discussed in the Supplementary Method 3). Therefore, acknowledging that their BP_ND_ estimates may be more affected by this non-equilibrium, the results involving this ROIs were still presented and interpreted in this manuscript. BP_ND_^0^ values were computed per session. Non-displaceable binding potentials (BP_ND_) were estimated by fitting a three-parameter non-linear Simplified Reference Tissue Model (SRTM)^86,87^ to the pre-task frames only, Using cerebellar cortex as the reference region. The cerebellar efflux constant k₂′ was fixed to the value previously estimated by multilinear reference tissue modelling^88^. This model is expressed by the following equation:

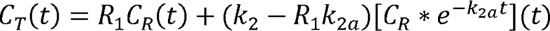

### PET Data Analysis Statistical Methods

BP_ND_^0^ data underwent outlier detection and imputation (see Supplementary Method 4 for details). Repeated-measures ANOVA with the factors Areas (nine ROIs) and Sessions (high-vs. low-reward) assessed session-related differences. As there was no session effect found for BP_ND_^0^, each session was treated as an independent data point for subsequent analyses. This approach allowed us to assess how an individual’s dopaminergic receptor availability might differentially relate to recognition memory across distinct reward contexts, despite reflecting the same underlying DA system health. These values were then correlated with corresponding HFA, controlling for age and applying FDR correction. This approach examined how interindividual differences in dopaminergic health (BP_ND_^0^) relate to memory performance.

### Interpretation of BP_ND_^0^

It has been reported that [^18^F]fallypride BP_ND_ reflects both D2/D3 receptor availability and competition from endogenous DA ^89,90^. In the absence of a task, BP_ND_^0^ is best understood as a composite measure that primarily indexes D2/D3 receptor density, and by extension, the health of the dopaminergic system, while still being partly and idiosyncratically influenced by baseline DA occupancy in each region. In source regions such as the SN, VTA, and LC, lower BP_ND_^0^ may indicate fewer unoccupied receptors or greater overall receptor loss, which could point to diminished inhibitory feedback or altered dopaminergic function ^28,71,91,92^. In terminal regions such as the thalamus, amygdala, and hippocampus, comparatively lower BP_ND_^0^ may likewise reflect reduced receptor availability, potentially linked to the region’s dopaminergic integrity. Viewed in this light, the measure is not a direct readout of ‘resting DA activity’ alone but rather a window onto how intact the D2/D3 receptor system is, and thus how well it might support episodic memory in older adults. We therefore interpret BP_ND_^0^ in older adults primarily as a proxy marker of dopaminergic system health, while recognising that some fraction of the signal arises from endogenous DA competition. Additional methodological details and supplementary analyses are provided in the Supplementary Method 4.

### Principal Axis Factoring of BP_ND_^0^

An exploratory factor analysis was performed on baseline D2/D3 receptor availability (BP_ND_^0^) to investigate whether nine ROIs could be reduced to a smaller number of latent factors^93^, using a custom R script with *psych* package (version 2.5.3.) and SPSS (SPSS v29.0.2.0). We focused on an averaged BP_ND_^0^ where high and low-reward values were averaged for each participant across the two sessions as there was no significant session difference (Figure 1). Prior to exploratory factor analysis, we evaluated sampling adequacy with KMO, tested whether the correlation matrices were factorable via Bartlett’s Test of Sphericity, and examined communalities for each ROI. Principal axis factoring was selected because our goal was to capture underlying shared variance among the ROIs while treating unique and error variance separately. A promax (oblique) rotation was used so that factors could be correlated. The number of factors to extract was determined by: (a) scree plot inspection, (b) Kaiser–Guttman criterion^94,95^, and (c) parallel analysis, which compares observed eigenvalues to those from randomly permuted data. All three criteria consistently supported a two-factor solution in each dataset (Supplementary Method 5 and Supplementary Figure 4).

### Volumetric Analysis in the ROIs

For each subject, first, intracranial volumes were calculated from a binary mask combining grey matter, white matter, and cerebrospinal fluid tissue classes estimated from their native whole-brain T1w images using SPM12’s Segment function. Afterwards, ROI volumes of the amygdala, caudate, NAcc, putamen, thalamus, SN, and VTA were calculated from the whole-brain parcellation generated from Freesurfer’s recon-all in their native space. Hippocampal volume was derived from each participant’s segmentation of aggregated and binarised hippocampal subfields using the Automatic Segmentation of Hippocampal Subfields (ASHS)^96^, followed by manual outline correction to improve anatomical accuracy. Normalised volumes for those ROIs were then computed using a custom MATLAB script according to the following formula:

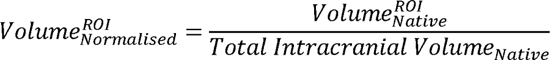

For the LC, canonical volumetric contrast was calculated from manual segmentations performed by an expert rater (YY) on session-averaged magnetisation transfer contrast (MTC) images. These averages were generated from four MTC acquisitions (two per scan session) using the *AverageImages* function from ANTs. Mean LC signal intensity was extracted from the labelled region and normalised against the mean intensity of a co-registered pons mask (transformed from MNI space to each subject’s native MTC space). The LC contrast ratio was computed as:

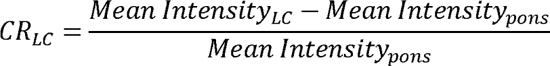

### Behavioural Analysis

For memory performance, *Hit-minus-FA* rates (Hit - FA; HFA) were used, maintaining interindividual variability rather than normalising with *D’*. Missing data for behavioural and other measures were handled via residual-based regression imputation (SPSS v29.0.2.0), excluding variables with >20% missing data as predictors. Further details on data handling of behavioural measures are described in the Supplementary Method 4.

## Authour contributions

YY, DH, OSp, and ED contributed to the conceptualisation and methodology of the study. YY contributed to investigation, analysis, visualisation, and writing of the original draft. JJ, JD, and CS provided analytical consultation. DH and ED supervised the investigation, analysis, and writing of the original draft. YY, MK, OSa, PS, KB, BG, MP, AS, SH, PG, SS, and IB contributed to the data collection. All authours reviewed and edited the manuscript.

## Declarations of interest

All authours report no conflict of interest.

## Notes

### Competing Interest Statement

The authors have declared no competing interest.

### Funding

- Deutsche Forschungsgemeinschaft: SFB1315 B06
- Deutsche Forschungsgemeinschaft: SFB1436 A08
- Deutsche Forschungsgemeinschaft: SFB779 A07
- AlzheimerM-BM-^Rs Research UK: SRF2018B-004

