## Supplementary Materials for "Different dopaminergic circuits defined by D2/D3 receptor availablity patterns show structure-specific links to memory in ageing"

**Abstract**

Cognitive ageing is marked by progressive decline in episodic memory and dopaminergic function, yet the extent to which individual differences in dopaminergic system integrity influence memory under motivational contexts remains unclear. In this study, we investigated how baseline D2/D3 receptor availability (BP_ND_^0^) in key dopaminergic pathways relates to reward-modulated memory performance in healthy older adults. Thirty-three healthy seniors (aged 64-85) underwent two session concurrent MR-PET imaging with [¹⁸F]fallypride involving scene categorisation task with high- and low-motivational contexts. We quantified BP_ND_^0^ across nine dopaminergic regions of interest and examined their relationships with recognition memory performance at short (~15m) and long (~24h) delays.

Baseline D2/D3 receptor availability showed high test-retest reliability and regionally distinct profiles, suggesting it reflects a stable neurochemical characteristic in healthy ageing, and principal axis factor analysis revealed two partially independent dopaminergic subsystems (dorsal striatal vs mesolimbic) based on interindividual patterns in receptor densities. Region-specific associations further linked D2/D3 receptor availability to distinct memory outcomes. Higher caudate D2/D3 receptor availability was associated with a liberal response bias (increased hits and false alarms both), whereas higher putamen D2/D3 predicted more durable long-term memory retention. Greater thalamic D2/D3 receptor availability correlated with fewer short-term false memories, while greater D2/D3 receptor availability in the amygdala was associated with better recognition at longer delays. In contrast, higher midbrain (substantia nigra and ventral tegmental area) D2/D3 availability which was linked to poorer reward-related memory performance. These findings suggest several complementary dopaminergic circuits supporting episodic memory. Our results highlight dopaminergic neuromodulation as a key factor in cognitive ageing and a potential target for interventions to bolster memory in late life.

**1. Supplementary method 1: Experimental protocols**

**For both visits:**

- Delayed memory test

**Visit 1 covariates:**

- Go-noGo task
- PSQI
- Raven’s matrices

**Visit 2 covariates:**

- CERAD
- BDI

**24-hour delay**

**Immediate memory test**

**Break**

**PET:**

- Baseline
- Task

**MRI:**

- 2^nd^ MPRAGE
- Fieldmap
- EPI

**PET:**

- In-Flow

**MRI:**

- 1^st^ MPRAGE
- MTC
- FLAIR
- T2 hippocampus
- T2 TSE
- QSM
- ASL

**Subject**

**preparation:**

*Changing clothes, collecting blood samples, placing flexure needle, performing two practice tasks*

radio-

activity

time🡪

**0m**

**95m**

**115m**

InFlow	Baseline	**Task**

**(High- or Low-Reward)**

**ca. 24h after the task**

Supplementary Figure 1. A detailed schedule of the experimental session. Subjects underwent two MR-PET scanning sessions, comprising high-reward and low-reward conditions, along with two post-scan covariate sessions 24 hours after each MR-PET scan. The sessions occurred on separate days, with an average inter-session interval of approximately 42 days. Experimental schedules were identical across sessions, except for the cognitive task performed inside the scanner, which had different reward conditions and stimulus sets based on a randomised list. Prior to each MR-PET session, subjects received task instructions and performed two practice tasks. Tracer administration and imaging procedures were conducted during the MR-PET scan. Following the scan, subjects underwent an immediate memory test, and subsequent post-scan covariate sessions included additional cognitive and psychological assessments as well as a delayed memory test.

All subjects underwent two MR-PET scanning sessons, consisting of a high-reward and a low-reward condition, as well as two post-scan covariate sessions 24 hours after each MR-PET scan. These four sessions took place on separate days, with a mean inter-session interval of 42.2±63.3 (*M*±*SD*) days. Each session had the same experimental schedule, except for the main cognitive task, which was performed inside the scanner and had different reward conditions and sets of stimuli depending on a predefined randomisation list. Upon arriving at the institute for each MR-PET session, subjects were once again instructed on the experimental tasks and informed of the schedule for both sessions. After instruction, they performed two 10-minute practice tasks on a computer to familiarise themselves to the button presses, task screens, and the task contingency and reward context of both scan sessions. Once the practice tasks were completed, subjects were moved to a preparation room, where two 5ml blood samples were taken in two 6ml EDTA tubes. They then waited in the preparation room until the tracer, which had been synthesized and delivered from the University Clinic Leipzig, arrived at the institute and passed quality assessment. The details of the synthesis procedure are described in the following section.

Once the tracer passed quality assessment procedure, subjects were equipped with Siemens pulse-oximetry and a breathing belt, positioned supine in the scanner, and underwent an MR-guided attenuation correction scan (MRAC). A physician then initiated intravenous bolus injection of the tracer 60 seconds after the first PET scan was started. The In-Flow PET scan, which was acquired to measure the radioactivity ramp-up, lasted for 60 minutes, during which subjects were instructed to stay awake. Concurrently with the PET scan, a series of structural MRI images were acquired.

After the first PET scan, subjects were given a 15-minute bathroom break outside the scanner, during which interaction with experimenters and physicians was controlled and minimised. Once they returned to the scanner, another MRAC was acquired, followed by a second 15-minute PET scan that measured baseline radioactivity levels. During the second PET scan, MRAC, a whole-brain T1-weighted image, and double-echo fieldmap images were acquired simultaneously. Following the scan, while inside the scanner, subjects were once again reminded and instructed on the reward task via an MR-compatible display positioned outside the bore and visible through a mirror attached to the head coil at eye level. 115 minutes after tracer injection, subjects began the task while the final 55-minute PET scan was performed. Once the task was completed, subjects were asked to rest with their eyes closed with an on-screen instruction while the remainder of the PET scan was conducted. After the scan, subjects' blood pressure was measured while they lay on the MRI table. They then performed an immediate recognition memory test (short-delay test) on a laptop for approximately 20 minutes in the preparation room. After the memory test, subjects were reminded of the post-scan covariate session scheduled for the next day and sent home.

Each of the two post-scan covariate sessions took place 24 hours after the subjects completed the reward task in the scanner. Upon arrival, they were briefly instructed on the second recognition memory test, which they performed on a desktop while their pupil diameter was measured. The contents of the two post-scan covariate sessions differed, except for the delayed incidental memory tests, which took place at the beginning of each session.

During the first post-scan session, after the delayed recognition memory test (long-delay test), subjects performed a 40-minute Go-noGo task on the same computer used for the memory test. Subjects then took a short 10-minute break and completed the C and D sets of Raven's matrices on paper, as well as the Pittsburgh Sleep Quality Index (PSQI). During the second post-scan session, subjects completed a full one-hour battery of the Consortium to Establish a Registry for Alzheimer's Disease (CERAD) and the Beck's Depression Inventory (BDI), both of which were paper-and-pencil assessments. A trained experimenter administered the CERAD test to the subjects.

**2. Supplementary Method 2: A full list of image acquisition parameters and PET reconstruction protocols**

In the first block of a scanning session, simultaneously with the first PET assessment, following structural MRI were acquired: Firstly, a MRAC was performed using a Dixon-based sequence to correct for attenuation of gamma rays in the PET reconstruction of three PET scans (voxel size = 1.3×1.3×2 mm, TE1 = 12.5 ms, TE2 = 25.1 ms, TR = 41 ms, flip angle [FA] = 10°, field of view [FOV] = 266×500×259 mm). A high-resolution T1-weighted magnetization-prepared rapid gradient echo (MPRAGE) was acquired for whole-brain segmentation that will be applied to PET data analysis and for guiding image registration (1 mm isotropic voxel size, 192 slices, TR = 2,500 ms, TE = 4.37 ms, TI = 1100ms, FOV = 256 mm, FA = 7°). A coronally oriented T2 image was acquired to capture hippocampus (voxel size = 0.4×0.4×2 mm, 29 slices, TR = 8020 ms, TE = 52 ms, FOV = 175 mm). In addition, an axially oriented T2-weighted fluid-attenuated inversion recovery (FLAIR) contrast was acquired (1 mm isotropic voxel size, 192 slices, TR = 5000 ms, TE = 396 ms, TI = 1800 ms, FOV = 256 mm). Afterwards, a T2-weighted turbo-spin echo image was acquired (0.5×0.5×2 mm, 80 slices, TR = 6910 ms, TE = 78 ms, FOV = 256 mm, FA = 120°). Following this, a quantitative susceptibility mapping (QSM) was acquired (0.5×0.5×1.4 mm, 104 slices, TR = 30 ms, TE = 15 ms, FOV = 224 mm, FA = 12°). Two gradient-echo (GRE) magnetization transfer contrasts (MTC) were acquired and later averaged together (0.8×0.8×1 mm, 64 slices, TR = 45 ms, TE = 5.36 ms, FOV = 200 mm, FA = 23°). Before the subjects went on a 15-minute break, two pulsed arterial spin labelling (pASL) with FAIR labelling and 3D-CREASE readout were acquired (2×2×5.5 mm, 30 slices, TR = 6000 ms, TE = 22.32 ms, BW = 2297 Hz/Px, EPI factor = 34, TF=10, two averaged multi-TI measurements with 13 TIs starting from 300 ms with an increment of 200 ms, FOV = 256 mm, FA = 120°).

In the second block of the scanning session after the break, another MRAC was acquired. Following MRAC, the second T1w image was acquired to assist functional MRI registration and PET data analysis as a source image for whole-brain segmentation (1 mm isotropic voxel size, 192 slices, TR = 2,500 ms, TE = 4.37 ms, TI = 1100ms, FOV = 256 mm, FA = 7°). Shortly before functional MRI acquisition, double-echo gradient echo fieldmap was acquired to guide the fMRI data unwarping (TR = 355ms, TE1 = 4.92 ms, TE2 = 7.38 ms, FA = 60°, FOV = 240×240×102 mm). Lastly, axially oriented T2*-weighted 2D EPIs with GRAPPA acceleration factor 2 were acquired during the reward task (2 mm isotropic voxel size, 51 slices, TR = 3600 ms, TE = 32 ms, FOV = 240×240×102 mm, FA = 80°).

The PET image acquisition involved using the radioligand [^18^F]fallypride (injected amount per subject: *M*±*SD*=193.73±3.80MBq, 0-175 minutes) to index dopamine D2D3R availability in striatum and extrastriatal regions. The precursor for [^18^F]fallypride synthesis was obtained from ABX (Radeberg, Germany) and Pharmasynth (Tartu, Estonia) respectively and labelling was performed on-site using a GE Tracerlab FX-FN synthesis module (GE Healthcare, Düsseldorf, Germany). The final product was obtained after reversed-phase high performance liquid chromatographic (HPLC) purification using initially a Supelco Ascentis C8 10 µm 250 mm x 10 mm column and 0.25 M ammonium acetate buffer pH 5/acetonitrile 70/30 v/v as mobile phase at a flow rate of 5 mL/min and later on a Supelcosil LC-8 column with a pore size of 5 µm and the dimensions 250 mm x 10 mm using the same eluent and flow. 18F-fallypride eluted after 8 min and 14 min respectively. The collected peak was diluted with 40 mL 0.15 M disodium hydrogen phosphate. The raw product was than fixated on a Waters Sep-Pak tC18 Plus Long solid phase extraction cartridge, washed with 10 mL water, eluted successively with 1 mL ethanol and 9 mL saline and sterile filtered over a Millipore Millex-GV 0.22 µm filter SLGVM33RS. The final product was buffered with 0.1 mL Natriumphosphat Braun (sodium phosphate, B. Braun Melsungen, Germany). The average specific radioactivity at the end of synthesis was > 8 GBq/mmol and the average radiochemical purity exceeded 97%.

The first PET scan consisted of dynamic data acquisition over 60 minutes (with 60 frames for 60 sec/frame), with this portion representing the ramp-up kinetics of the [^18^F]fallypride. After this scan, subjects had a 15-minute break outside the scanner. The second PET scan involved repositioning subjects in the scanner, acquiring a 2-minute magnetic Resonance Attenuation Correction (MRAC) scan, and acquiring the second dynamic PET scan for 15 minutes (15 frames for 60 sec/frame), which served as the baseline radioactivity reference scan. The third and final dynamic PET scan, during which subjects performed a cognitive task, was acquired for 55 minutes (55 frames for 60 sec/frame). All PET scans were acquired with a voxel size of 2.3×2.3×5 mm.

For the PET image reconstruction protocol, the following specifications were adhered to across three separate acquisitions. The imaging was performed using a high-definition PET (HD-PET) algorithm with a set number of three iterations and an image resolution of 256x256 mm. Images were magnified using a zoom factor of 3.0, and a Gaussian filter with a kernel size of 2.0 was applied for smoothing. Additionally, scatter correction was consistently employed for all scans.

**3. Supplementary Method 3: Image data preprocessing and analysis procedure**

**Structural whole-brain (T1w) data**: Two T1w images of each dataset were first bias-field corrected and segmented for grey matter, white matter, and CSF using SPM12’s *Segment* function. During this step, the default bias-field correction parameters set by SPM12 were used besides *Bias FWHM (full width at half maximum)* option, which was set at 30mm in order to refine the bias-field inhomogeneity correction near cerebellar cortex, which caused under-segmentation of cerebellar cortex during the whole-brain parcellation. The bias-field corrected T1w images were then segmented whole-brain using Freesurfer’s *recon-all* function with *-all* switch, which enables the function to perform full routine of cortical reconstruction (version 7.4.1; Dale et al., 1998). The resulting whole-brain segmentation was rigidly registered back to respective native T1w image spaces from Freesurfer’s atlas space. Finally, every participant’s segmentation was visually inspected in native T1 space. Any inaccuracies in outlines of cerebellar cortex were corrected manually to ensure anatomical fidelity and avoid errors in the pharmacokinetic modelling associated with reference region.

As the next step, SN and VTA segmentations were extracted from a probabilistic subcortical brain nuclei atlas by Pauli, Nili and Tyszka (Pauli, Nili, and Tyszka 2018) using *fslmaths* of FSL toolbox (FMRIB Software Library; Jenkinson et al., 2012), and LC segmentation were taken from a biologically plausible volume of interest that shows high overlap across studies that examined LC (Dahl et al. 2022). These segmentations in each of their structural template space were non-linearly “**back-transformed**” onto each subject’s both of native T1w spaces using *antsRegistrationSyN* and *antsApplyTransforms* functions with nearest neighbour interpolation for the latter function. To enhance the precision of transformation study-specific structural template was used as an intermediate step. That is, [1] an individual whole-brain structural image (a) was first non-linearly registered to the study-specific template (b), and [2] the study-specific template was non-linearly registered to the structural MNI templates (Fonov et al. 2011) (c) SN, VTA, and LC atlases were delineated. Afterwards, the transformation matrices and inverse warps acquired from the previous non-linear transformation steps were concatenated to transform the atlases into each of subjects’ native geometrical spaces. More commonly, the order of those steps is inversed, moving the structural MNI images (c) into the native T1w images (a) directly. However, in our study, as a part of fMRI image preprocessing pipeline, step [1] and step [2] were already performed. Therefore, in order to save computational resources, the transformation steps for those atlases were adapted accordingly. The overview of this spatial transformation procedure is delineated in Supplementary Figure 2. The transformed SN, VTA, and LC atlases in each subject’s native space were then added to the existing whole-brain segmentations in native T1w image spaces acquired from the previous step using a custom MATLAB script.

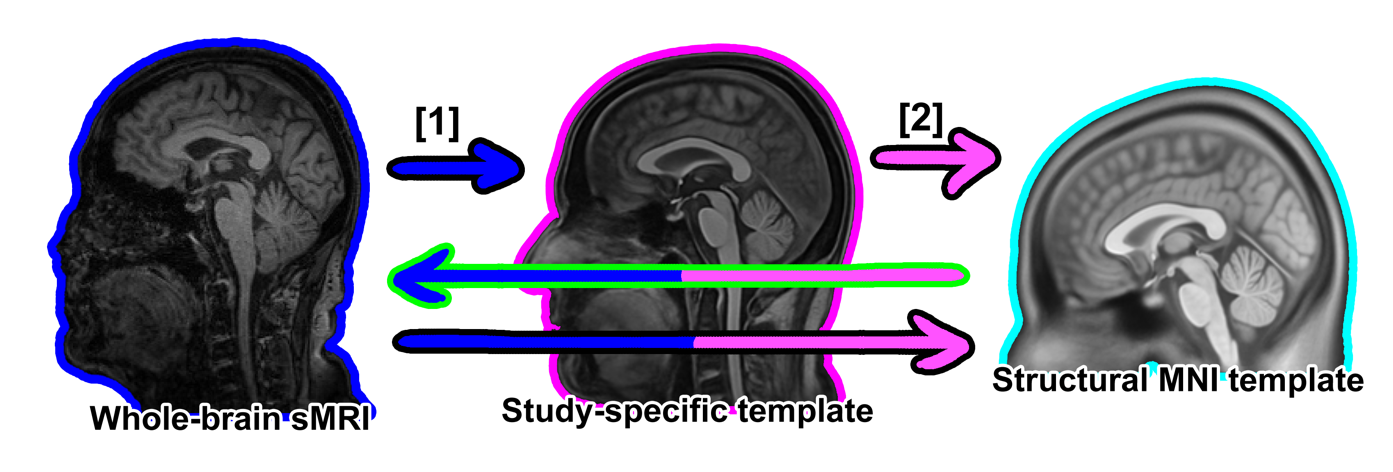

**(b)**

**(a)**

**(c)**

**Supplementary Figure 2. Overview of the structural MRI image co-registration steps.** An individual whole-brain structural image (a) was initially aligned non-linearly to a study-specific template (b) generated from each subject’s bias-field-corrected structural whole-brain images (indicated as a blue arrow). This template (b) was then non-linearly registered to the structural MNI templates (c) on which the SN, VTA, and LC atlases were delineated (indicated as a pink arrow). Subsequently, the transformation matrices and inverse warps derived from these registration steps were combined to ‘back-transform’ the atlases into each subject’s native anatomical space (indicated as a multicoloured arrow with green outline).

**PET data:** To estimate regional D2D3R binding, whole-brain segmentations acquired from two T1w images were used, which were augmented with subcortical segmentation transformed from subcortical atlases. Each PET scan’s frames were registered rigidly with linear interpolation to the native structural T1w image of each scanning block they were concurrently scanned to, i.e., the first T1w image for in-flow mean PET scan and the second T1w scan of baseline and task mean PET scan, using the *antsRegistrationSyN.sh* function of ANTs (version 2.3.1; Avants et al., 2011).

The PET scans in each of respective native T1w space were then used as an input to a custom MATLAB script generating ROI-specific time-activity curves (TACs), which were later used as inputs to a custom MATLAB script performing pharmacokinetic modelling described in 5.1.4.2.

Regional baseline non-displaceable binding potential (*BP_ND_^0^*) values were obtained by fitting the three-parameter non-linear Simplified Reference Tissue Model (SRTM; Lammertsma & Hume, 1996) to the pre-task frames only, excluding all task phase data. Cerebellar cortex TACs served as the reference input. The SRTM is based on a two-tissue compartment model that estimates the binding potential without the need for arterial sampling. This model is expressed by the following equation:

$$C_{T}\left( t \right)=R_{1}C_{R}\left( t \right)+(k_{2}-R_{1}k_{2a}){[C}_{R}*e^{-k_{2a}t}](t)$$

In this equation, $C_{T}$ and $C_{R}$ denote the tracer concentrations in the target and reference regions, respectively. $R_{1}$ is the delivery ratio, reflecting the ratio of tracer delivery between the target and reference tissues; $k_{2}$ is the rate constant for the tracer's efflux from the target region to plasma; $k_{2a}$ is the apparent efflux constant when the reference input replaces the arterial input; and $t$ is the time in minutes. In our analysis, the cerebellar efflux constant k₂′ was fixed to the value previously estimated by multilinear reference tissue modelling (Ichise et al. 2003) and then held fixed during the non-linear SRTM fit to the baseline data. This model provides a robust framework for estimating receptor availability in the target regions relative to a non-displaceable reference region.

A quantitative measure of the non-displaceable binding potential (*BP_ND_*), representing receptor availability, is derived from the kinetic parameters of the model and is computed as follows:

$${BP}_{ND}=\frac{k_{2}}{k_{2a}}-1$$

The selection of ROIs for the PET component of the study was guided by a targeted approach to investigate the ageing neuromodulatory system and its impact on memory. The primary focus was on areas known to be rich in dopamine D2 and D3 receptors. To identify these regions, the Human Protein Atlas, which provides detailed expression profiles of various receptors throughout the brain, was consulted (Karlsson et al. 2021; Sjöstedt et al. 2020; Thul et al. 2017; Uhlen et al. 2017, 2019; Uhlén et al. 2015, 2019). However, not all regions displaying high receptor density were included as ROIs in the final binding potential analysis. The selection was refined to include only those areas that are specifically relevant to the study's aim of exploring how the ageing dopaminergic system affect episodic memory. This selective approach ensured that the ROIs were not only rich in the receptors of interest but also pertinent to the cognitive functions and age-related changes under the study’s investigative goals. By focusing on these particular regions, the study aims to yield more precise interpretations into the specific roles that dopaminergic activities play in memory processes in the ageing brain. A list of initially selected ROIs and their normalised receptor density is outlined in Table 3A.

|  | **Density (nTPM)** | | |  | $\frac{\boldsymbol{nTPM}_{\boldsymbol{D}\boldsymbol{1}}}{\boldsymbol{nTPM}_{\boldsymbol{D}\boldsymbol{2}\boldsymbol{D}\boldsymbol{3}}}$ |
| --- | --- | --- | --- | --- | --- |
| **ROI** | **DRD2** | **DRD3** | **Sum (D2+D3)** | **DRD1** |  |
| **Putamen** | 163.50 | 4.4 | **167.90** | 46.6 | 0.278 |
| **Caudate** | 113.20 | 4 | **117.20** | 44.5 | 0.379 |
| **Nucleus Accumbens** | 105.70 | 9.6 | **115.30** | 43.4 | 0.376 |
| **Substantia Nigra** | 88.60 | 0.4 | **89.00** | 0.4 | 0.004 |
| **Ventral Tegmental Area** | 61.20 | 0.8 | **62.00** | 0.6 | 0.010 |
| **Locus Coeruleus** | 46.60 | 0.6 | **47.20** | 0.6 | 0.013 |
| **Thalamus** | 32.80 | 0.8 | **33.60** | 4.1 | 0.122 |
| **Amygdala** | 7.60 | 0.7 | **8.30** | 6.8 | 0.819 |
| **Hippocampus** | 3.90 | 0.8 | **4.70** | 3.7 | 0.787 |

Supplementary Table 1. A list of selected ROIs included in the PET data analysis and their receptor density in human brain in normalized transcripts in million (nTPM).

From this initial list of ROIs, additional analyses were performed on these initial ROIs to ensure analysis validity and interpretability of BP measure. To verify whether the data had reached equilibrium, thereby allowing us to reliably capture the stability of the BP_ND_^0^, the slopes of the target-to-reference ratio curves were calculated across all subjects. These curves were smoothed using the locally estimated scatterplot smoothing method (LOESS), and linear fits were applied to compute the slopes of the target-to-reference curves from the decay-corrected baseline to task TACs. These slopes were then tested against those of the reference region, the cerebellar cortex, using a one-sample *t*-test.

Our analyses indicated that although TACs for the putamen, caudate, and nucleus accumbens levelled off, they did not achieve target-to-reference ratio equilibrium, showing the continued increase throughout the 15-min baseline scan (95-110 min after the tracer injection) and during the subsequent 55-min task scan (115-170 min; Supplementary Figure 3 and Supplementary Table 2). This indicates that BP_ND_^0^ values in these regions may be underestimated. This may be due to differences in equilibrium rates between areas with high and low receptor densities, as demonstrated by Ceccarini et al. (2012) in their simulations (Ceccarini et al. 2012; Christian et al. 2000). Although the putamen, caudate, and NAcc did not reach kinetic equilibrium in our scans (Ceccarini et al. 2012; Slifstein and Laruelle 2000), their baseline BP_ND_ (BP_ND_^0^) can still offer insight into overall receptor availability. Simulations and empirical work with [¹⁸F]fallypride show that reducing an 180-min [¹⁸F]fallypride scan *step-wise* by up to 60 min lowered BP_ND_ estimated with SRTM in the putamen by only ≈0.58% per ten-minute block (≤6% for a 60-min truncation), and, crucially, left between-subject rank order virtually unchanged (Vernaleken et al. 2011). Therefore, absolute values may be biased downward, but the individual-difference signal we correlate with behaviour is largely preserved.

On the other hand, two extrastriatal regions, the amygdala and hippocampus, showed pseudo-equilibrium. Although statistically significant increase was observed in the hippocampus and amygdala’s target-to-reference ratio (Supplementary Table 2 and Supplementary Figure 3), the magnitude of these changes, *M*±*SD*=0.0003±0.00001% for the amygdala and 0.0034±0.0016% for the hippocampus across all baseline and task PET scan timeline, which is 75-minute period. This means, the absolute change in target-to-reference ratios of 0.0018 and 0.0006 are below 0.2% of baseline signal. Simulation and test-retest studies place the extrastriatal detection limit for [^18^F]fallypride near 10% (Ceccarini et al. 2012; Christian et al. 2000), while Monte-Carlo work shows mean-zero counting noise alone can depress V_T_/BP_ND_ by up to 13% in hippocampus-sized VOIs (Slifstein and Laruelle 2000). The below 1% hippocampal and amygdalar slopes therefore fall inside expected error and are unlikely to reflect genuine neurobiological change.

Given these considerations, the observed increases in the target-to-reference ratio in the amygdala and hippocampus, despite being statistically significant, may not have practical significance due to the potential for noise and variability in these measurements. In addition, it is important to acknowledge that other physiological processes, such as receptor dynamics or metabolic fluctuations (Halldin et al. 1998), could contribute to these observed changes. For these reasons, while acknowledging that their BP_ND_ estimates may be more affected by this non-equilibrium, the results involving this ROIs were still presented and interpreted in this manuscript.

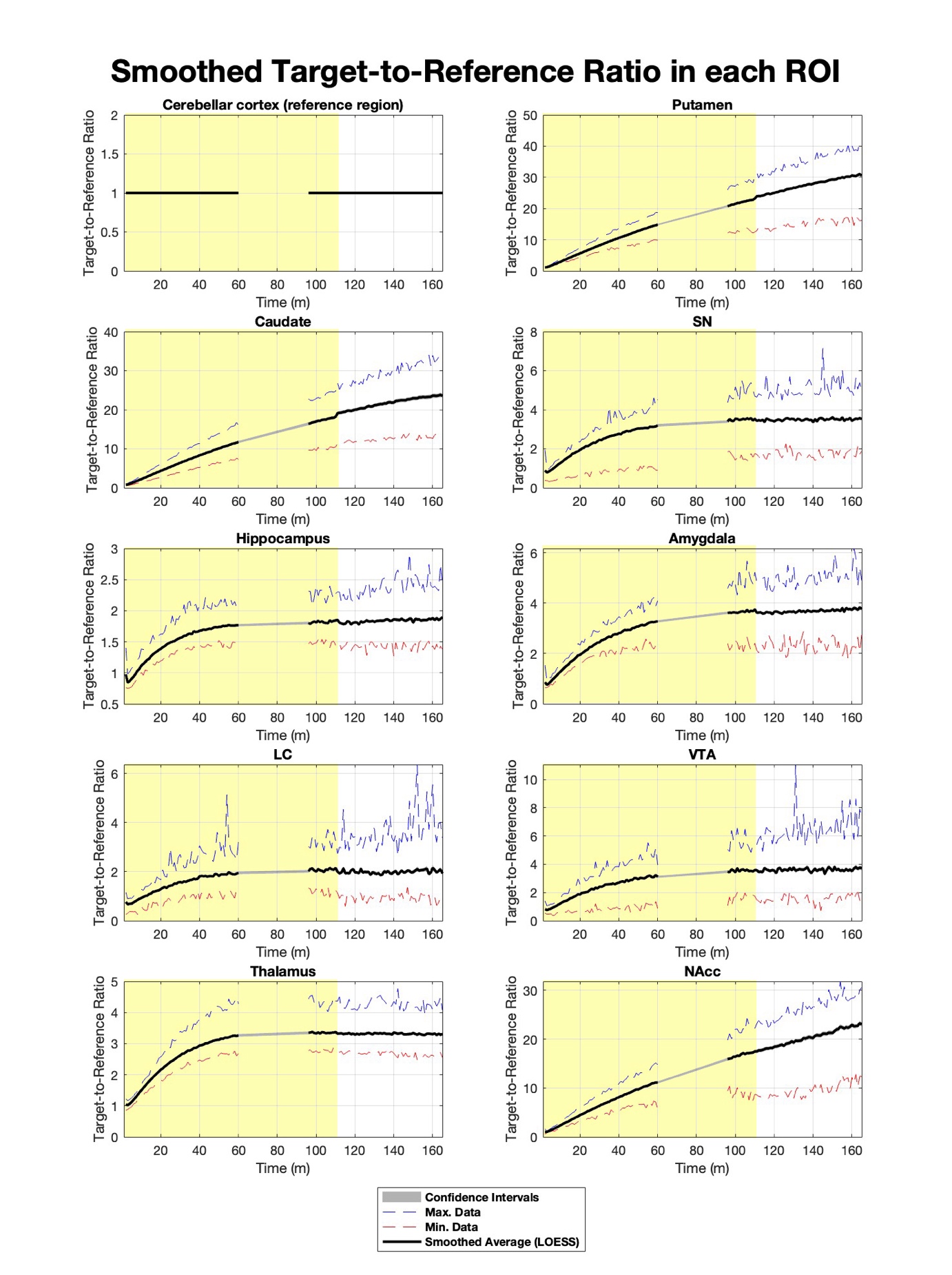

Supplementary Figure 3. LOESS-smoothed target-to-reference curves calculated from TACs from all subjects in both PET scan sessions.

This figure illustrates LOESS-smoothed target-to-reference ratio curves derived from TACs aggregated from all subjects during both PET scan sessions. Each plot represents a different study ROIs with the cerebellar cortex serving as the reference region. The solid black line indicates the smoothed average of the target-to-reference ratio, with dashed red and blue lines representing the maximum and minimum data points observed, respectively. The grey shaded areas around the solid black line represent the standard error around the smoothed average. The yellow shaded area in each plot indicates the time window (0 to 110 minutes post tracer injection) used to calculate BP_ND_^0^. The plots surrounded by red outlines shows the ROIs that were included in the final analyses. Statistical details of the slope computed within the baseline and the task scans are described in **Supplementary Table 2** below.

| **ROI** | **Slope (Linear)** | ***T*-statistics (df)** | **95% Confidence Interval**  **[lower bound, upper bound]** |
| --- | --- | --- | --- |
| **Putamen** | 0.00230 | 36.83 (59)* | [0.00210, 0.00250] |
| **Caudate** | 0.00169 | 27.55 (59)* | [0.00150, 0.00180] |
| **Nucleus Accumbens** | 0.00161 | 26.96 (59)* | [0.00140, 0.00180] |
| **Substantia Nigra** | ≅0.00001 | 1.08 (59) | [-0.00001, 0.00003] |
| **Ventral Tegmental Area** | 0.00003 | 2.35 (59) | [≅-0.00001, 0.00006] |
| **Locus Coeruleus** | ≅-0.00001 | -1.24 (59) | [-0.00003, 0.00001] |
| **Thalamus** | -0.00001 | -2.56 (59) | [-0.00002, 0.00000] |
| **Amygdala** | 0.00003 | 4.49* (59) | [0.00001, 0.00005] |
| **Hippocampus** | 0.00001 | 5.22* (59) | [0.00001, 0.00002] |

Supplementary Table 2. Statistical details of the target-to-reference ratio curve.

This table presents the statistical details of the LOESS-smoothed target-to-reference ratio curves for provisional study ROIs derived from TACs of all subjects across both PET scan sessions. The slope (calculated as linear fit) represents the gradient of the smoothed curves, indicating the rate of change over time. *T*-statistics are calculated from a one-sample t-test comparing each ROI’s target-to-reference ratio curve against the reference region’s target-to-reference ratio curve, treating each scan session as a distinct data point, and the values within the brackets represent degrees of freedom for each test. The 95% confidence interval is provided for the slope, detailing the estimated precision of the slope values. Asterisks (*) next to T-statistics signify statistical significance of *p*<.05.

**4. Supplementary Method 4: Data handling procedure and imputation statistics**

The data processing steps included removal of outliers and imputation for all measures presented in this manuscript.

In the dataset, memory test performance and BP_ND_^0^ were screened for outliers using the interquartile-range (IQR) method or Tukey's box-plot method, which is used to identify outliers by defining bounds as 1.5 times the interquartile range (IQR) below the first quartile (Q1) and above the third quartile (Q3) denoted in the following formula:

$$Lower bound=Q1-1.5\times IQR$$

$$Upper bound=Q3+1.5\times IQR$$

Afterwards, missing data patterns were assessed separately for BP_ND_^0^ and memory datasets in order to respect the contractual and distributional distinctions between neurochemical (BP_ND_^0^) and cognitive (memory) variables. Variables with more than 20% missing data were excluded from the predictor list for subsequent regression analyses to minimise potential biases associated with high levels of imputation and data sparsity (Supplementary Table 3). This threshold aligns with established guidelines in neuroimaging and clinical research literature, where a 20% cutoff is commonly used to balance data retention and validity of model estimation (Grassi et al. 2019; Shishegar et al. 2021). For included predictor variables (i.e., those with ≤20% missingness), missing values were handled using the regression method available in SPSS Missing Value Analysis (MVA; SPSS v29.0.2.0). This method estimates missing entries based on linear regression models using all other observed variables as predictors, assuming a missing-at-random mechanism. Regression imputation is particularly suitable when data points are missing at random pattern and when predictors are highly correlated with each other in the dataset (Azur et al. 2011; McCombe et al. 2022; Schafer and Graham 2002). Indeed, Little’s MCAR test yielded no significant result, indicating that the missing data of this dataset is likely random (Little 1988), *χ*^2^(435)=356.230, *p*=.998. All imputed values were inspected to ensure consistency with observed distributions (Supplementary Table 4), and no outliers beyond 1.5×IQR were introduced through this process.

| Session | Variable | N | Missing | Mean | SD | % Missing | Excluded from predictor list (Missing > 20%) |
| --- | --- | --- | --- | --- | --- | --- | --- |
| High Reward | BP_ND_^0^ Amydala | 28 | 5 | 2.069 | 0.203 | 15.2 | FALSE |
|  | BP_ND_^0^ Caudate | 30 | 3 | 19.393 | 3.522 | 9.1 | FALSE |
|  | BP_ND_^0^ Hippocampus | 29 | 4 | 0.610 | 0.099 | 12.1 | FALSE |
|  | BP_ND_^0^ LC | 30 | 3 | 0.806 | 0.145 | 9.1 | FALSE |
|  | **BP_ND_^0^ NAcc** | **25** | **8** | **19.904** | **2.621** | **24.2** | **TRUE** |
|  | BP_ND_^0^ Putamen | 27 | 6 | 25.078 | 2.543 | 18.2 | FALSE |
|  | BP_ND_^0^ SN | 29 | 4 | 1.951 | 0.223 | 12.1 | FALSE |
|  | BP_ND_^0^ Thalamus | 29 | 4 | 1.851 | 0.192 | 12.1 | FALSE |
|  | BP_ND_^0^ VTA | 30 | 3 | 2.041 | 0.377 | 9.1 | FALSE |
| Low Reward | BP_ND_^0^ Amygdala | 30 | 3 | 2.083 | 0.236 | 9.1 | FALSE |
|  | BP_ND_^0^ Caudate | 27 | 6 | 20.166 | 2.640 | 18.2 | FALSE |
|  | BP_ND_^0^ Hippocampus | 30 | 3 | 0.621 | 0.108 | 9.1 | FALSE |
|  | BP_ND_^0^ LC | 29 | 4 | 0.762 | 0.159 | 12.1 | FALSE |
|  | BP_ND_^0^ NAcc | 30 | 3 | 19.444 | 3.238 | 9.1 | FALSE |
|  | BP_ND_^0^ Putamen | 29 | 4 | 25.529 | 3.029 | 12.1 | FALSE |
|  | BP_ND_^0^ SN | 28 | 5 | 1.941 | 0.218 | 15.2 | FALSE |
|  | BP_ND_^0^ Thalamus | 29 | 4 | 1.854 | 0.220 | 12.1 | FALSE |
|  | BP_ND_^0^ VTA | 29 | 4 | 1.980 | 0.354 | 12.1 | FALSE |
| High Reward | HFA All Scenes_Short-delay | 30 | 3 | 0.351 | 0.221 | 9.1 | FALSE |
|  | HFA All Scenes Long-delay | 28 | 5 | 0.325 | 0.211 | 15.2 | FALSE |
|  | HFA Reward Short-delay | 30 | 3 | 0.327 | 0.245 | 9.1 | FALSE |
|  | HFA Reward Long-delay | 28 | 5 | 0.314 | 0.267 | 15.2 | FALSE |
|  | HFA Neutral Short-delay | 30 | 3 | 0.361 | 0.265 | 9.1 | FALSE |
|  | HFA Neutral Long-delay | 28 | 5 | 0.306 | 0.237 | 15.2 | FALSE |
|  | Hits All Scenes Short-delay | 30 | 3 | 0.808 | 0.147 | 9.1 | FALSE |
|  | Hits All Scenes Long-delay | 28 | 5 | 0.741 | 0.22 | 15.2 | FALSE |
|  | Hits Reward Short-delay | 28 | 5 | 0.847 | 0.121 | 15.2 | FALSE |
|  | Hits Reward Long-delay | 28 | 5 | 0.758 | 0.242 | 15.2 | FALSE |
|  | Hits Neutral Short-delay | 30 | 3 | 0.791 | 0.171 | 9.1 | FALSE |
|  | Hits Neutral Long-delay | 28 | 5 | 0.716 | 0.255 | 15.2 | FALSE |
|  | FA All Scenes Short-delay | 30 | 3 | 0.457 | 0.268 | 9.1 | FALSE |
|  | FA All Scenes Long-delay | 28 | 5 | 0.416 | 0.310 | 15.2 | FALSE |
|  | FA Reward Short-delay | 30 | 3 | 0.495 | 0.281 | 9.1 | FALSE |
|  | FA Reward Long-delay | 28 | 5 | 0.443 | 0.346 | 15.2 | FALSE |
|  | FA Neutral Short-delay | 30 | 3 | 0.43 | 0.3 | 9.1 | FALSE |
|  | FA Neutral Long-delay | 28 | 5 | 0.410 | 0.334 | 15.2 | FALSE |
| Low Reward | HFA All Scenes_Short-delay | 29 | 4 | 0.332 | 0.209 | 12.1 | FALSE |
|  | HFA All Scenes Long-delay | 28 | 5 | 0.333 | 0.242 | 15.2 | FALSE |
|  | HFA Reward Short-delay | 29 | 4 | 0.265 | 0.245 | 12.1 | FALSE |
|  | HFA Reward Long-delay | 28 | 5 | 0.301 | 0.271 | 15.2 | FALSE |
|  | HFA Neutral Short-delay | 29 | 4 | 0.372 | 0.269 | 12.1 | FALSE |
|  | **HFA Neutral Long-delay** | **26** | **7** | **0.286** | **0.215** | **21.2** | **TRUE** |
|  | Hits All Scenes Short-delay | 28 | 5 | 0.831 | 0.13 | 15.2 | FALSE |
|  | Hits All Scenes Long-delay | 27 | 6 | 0.79 | 0.161 | 18.2 | FALSE |
|  | Hits Reward Short-delay | 28 | 5 | 0.819 | 0.142 | 15.2 | FALSE |
|  | Hits Reward Long-delay | 27 | 6 | 0.807 | 0.166 | 18.2 | FALSE |
|  | Hits Neutral Short-delay | 29 | 4 | 0.827 | 0.161 | 12.1 | FALSE |
|  | Hits Neutral Long-delay | 27 | 6 | 0.77 | 0.173 | 18.2 | FALSE |
|  | FA All Scenes Short-delay | 29 | 4 | 0.483 | 0.236 | 12.1 | FALSE |
|  | FA All Scenes Long-delay | 28 | 5 | 0.429 | 0.260 | 15.2 | FALSE |
|  | FA Reward Short-delay | 29 | 4 | 0.538 | 0.269 | 12.1 | FALSE |
|  | FA Reward Long-delay | 28 | 5 | 0.476 | 0.312 | 15.2 | FALSE |
|  | FA Neutral Short-delay | 29 | 4 | 0.454 | 0.270 | 12.1 | FALSE |
|  | FA Neutral Long-delay | 28 | 5 | 0.408 | 0.277 | 15.2 | FALSE |

**Supplementary Table 3. A table outlining the pre-imputation descriptive statistics and percentage of missing data of all variables. The last column indicates which variable has been excluded from the final data imputation.**

|  |  | **Pre-Imputation** | | |  | **Post-Imputation** | | |
| --- | --- | --- | --- | --- | --- | --- | --- | --- |
| **Session** | **Variable** | **Mean** | **SD** | **Variance** |  | **Mean** | **SD** | **Variance** |
| High Reward | BP_ND_^0^ Amydala | 2.092 | 0.166 | 0.041 |  | 2.062 | 0.190 | 0.036 |
|  | BP_ND_^0^ Caudate | 20.639 | 3.178 | 12.405 |  | 19.438 | 3.361 | 11.297 |
|  | BP_ND_^0^ Hippocampus | 0.652 | 0.067 | 0.01 |  | 0.604 | 0.097 | 0.009 |
|  | BP_ND_^0^ LC | 0.807 | 0.111 | 0.021 |  | 0.806 | 0.140 | 0.02 |
|  | BP_ND_^0^ NAcc | 19.401 | 2.606 | 6.869 |  | 19.977 | 2.522 | 6.363 |
|  | BP_ND_^0^ Putamen | 24.810 | 2.813 | 6.468 |  | 24.999 | 2.590 | 6.706 |
|  | BP_ND_^0^ SN | 1.967 | 0.171 | 0.05 |  | 1.954 | 0.209 | 0.044 |
|  | BP_ND_^0^ Thalamus | 1.872 | 0.163 | 0.037 |  | 1.852 | 0.184 | 0.034 |
|  | BP_ND_^0^ VTA | 2.130 | 0.283 | 0.142 |  | 2.057 | 0.370 | 0.137 |
| Low Reward | BP_ND_^0^ Amygdala | 2.089 | 0.211 | 0.056 |  | 2.082 | 0.232 | 0.054 |
|  | BP_ND_^0^ Caudate | 20.735 | 2.129 | 6.972 |  | 19.896 | 2.509 | 6.296 |
|  | BP_ND_^0^ Hippocampus | 0.647 | 0.091 | 0.012 |  | 0.620 | 0.104 | 0.011 |
|  | BP_ND_^0^ LC | 0.750 | 0.147 | 0.025 |  | 0.773 | 0.156 | 0.024 |
|  | BP_ND_^0^ NAcc | 18.962 | 2.792 | 10.484 |  | 19.405 | 3.143 | 9.879 |
|  | BP_ND_^0^ Putamen | 25.084 | 2.673 | 9.176 |  | 25.531 | 2.984 | 8.906 |
|  | BP_ND_^0^ SN | 1.951 | 0.213 | 0.048 |  | 1.945 | 0.203 | 0.041 |
|  | BP_ND_^0^ Thalamus | 1.868 | 0.157 | 0.049 |  | 1.859 | 0.211 | 0.044 |
|  | BP_ND_^0^ VTA | 1.936 | 0.335 | 0.125 |  | 1.985 | 0.332 | 0.11 |
| High Reward | HFA All Scenes_Short-delay | 0.356 | 0.214 | 0.046 |  | 0.356 | 0.214 | 0.046 |
|  | HFA All Scenes Long-delay | 0.325 | 0.196 | 0.039 |  | 0.325 | 0.196 | 0.039 |
|  | HFA Reward Short-delay | 0.330 | 0.241 | 0.058 |  | 0.330 | 0.241 | 0.058 |
|  | HFA Reward Long-delay | 0.322 | 0.247 | 0.061 |  | 0.322 | 0.247 | 0.061 |
|  | HFA Neutral Short-delay | 0.359 | 0.254 | 0.065 |  | 0.359 | 0.254 | 0.065 |
|  | HFA Neutral Long-delay | 0.299 | 0.221 | 0.049 |  | 0.299 | 0.221 | 0.049 |
|  | Hits All Scenes Short-delay | 0.801 | 0.147 | 0.022 |  | 0.801 | 0.147 | 0.022 |
|  | Hits All Scenes Long-delay | 0.742 | 0.207 | 0.043 |  | 0.742 | 0.207 | 0.043 |
|  | Hits Reward Short-delay | 0.835 | 0.117 | 0.014 |  | 0.835 | 0.117 | 0.014 |
|  | Hits Reward Long-delay | 0.736 | 0.234 | 0.055 |  | 0.736 | 0.234 | 0.055 |
|  | Hits Neutral Short-delay | 0.787 | 0.164 | 0.027 |  | 0.787 | 0.164 | 0.027 |
|  | Hits Neutral Long-delay | 0.735 | 0.244 | 0.059 |  | 0.735 | 0.244 | 0.059 |
|  | FA All Scenes Short-delay | 0.446 | 0.262 | 0.068 |  | 0.446 | 0.262 | 0.068 |
|  | FA All Scenes Long-delay | 0.425 | 0.288 | 0.083 |  | 0.425 | 0.288 | 0.083 |
|  | FA Reward Short-delay | 0.501 | 0.272 | 0.074 |  | 0.501 | 0.272 | 0.074 |
|  | FA Reward Long-delay | 0.452 | 0.322 | 0.103 |  | 0.452 | 0.322 | 0.103 |
|  | FA Neutral Short-delay | 0.424 | 0.287 | 0.082 |  | 0.424 | 0.287 | 0.082 |
|  | FA Neutral Long-delay | 0.412 | 0.329 | 0.108 |  | 0.412 | 0.329 | 0.108 |
| Low Reward | HFA All Scenes_Short-delay | 0.315 | 0.204 | 0.041 |  | 0.315 | 0.204 | 0.041 |
|  | HFA All Scenes Long-delay | 0.320 | 0.234 | 0.055 |  | 0.320 | 0.234 | 0.055 |
|  | HFA Reward Short-delay | 0.270 | 0.238 | 0.057 |  | 0.270 | 0.238 | 0.057 |
|  | HFA Reward Long-delay | 0.325 | 0.284 | 0.081 |  | 0.325 | 0.284 | 0.081 |
|  | HFA Neutral Short-delay | 0.388 | 0.263 | 0.069 |  | 0.388 | 0.263 | 0.069 |
|  | HFA Neutral Long-delay | 0.273 | 0.209 | 0.044 |  | 0.273 | 0.209 | 0.044 |
|  | Hits All Scenes Short-delay | 0.824 | 0.126 | 0.016 |  | 0.824 | 0.126 | 0.016 |
|  | Hits All Scenes Long-delay | 0.795 | 0.149 | 0.022 |  | 0.795 | 0.149 | 0.022 |
|  | Hits Reward Short-delay | 0.804 | 0.151 | 0.023 |  | 0.804 | 0.151 | 0.023 |
|  | Hits Reward Long-delay | 0.810 | 0.154 | 0.024 |  | 0.810 | 0.154 | 0.024 |
|  | Hits Neutral Short-delay | 0.818 | 0.155 | 0.024 |  | 0.818 | 0.155 | 0.024 |
|  | Hits Neutral Long-delay | 0.778 | 0.175 | 0.031 |  | 0.778 | 0.175 | 0.031 |
|  | FA All Scenes Short-delay | 0.494 | 0.228 | 0.052 |  | 0.494 | 0.228 | 0.052 |
|  | FA All Scenes Long-delay | 0.421 | 0.245 | 0.060 |  | 0.421 | 0.245 | 0.060 |
|  | FA Reward Short-delay | 0.544 | 0.258 | 0.066 |  | 0.544 | 0.258 | 0.066 |
|  | FA Reward Long-delay | 0.480 | 0.298 | 0.089 |  | 0.480 | 0.298 | 0.089 |
|  | FA Neutral Short-delay | 0.449 | 0.258 | 0.067 |  | 0.449 | 0.258 | 0.067 |
|  | FA Neutral Long-delay | 0.390 | 0.267 | 0.071 |  | 0.390 | 0.267 | 0.071 |

**Supplementary Table 4. A table comparing the pre- and post-imputation descriptive statistics.** No variable has shown a significant change in their descriptive statistics after the data imputation.

**5. Supplementary Method 5: Detailed diagnostics performed for principal axis factoring (PAF) analysis performed on the BP_ND_^0^ values averaged across the two sessions**

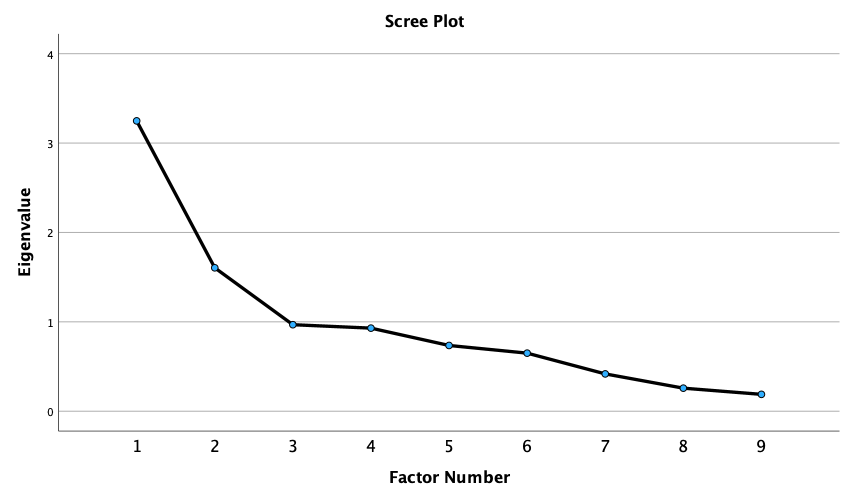
**Supplementary Figure 4.** **Scree plot of the nine eigenvalues from the PAF analysis on BP_ND_^0^** **values.**

An exploratory principal‐axis factor analysis was conducted on BP_ND_^0^ values averaged across two sessions from nine ROIs. Following recommended practice (Costello and Osborne 2005; Gorsuch 1973), factorability was first assessed: the Kaiser-Meyer-Olkin measure of sampling adequacy was .588 and Bartlett’s test of sphericity was significant, *χ*²(36) = 86.99, *p*<.001, indicating the data matrix was appropriate for factoring analysis. Initial eigenvalues from the unrotated solution yielded two factors over eigenvalue of 1 (Factor 1=3.248; Factor 2=1.604), explaining 36.1% and 17.8% of the total variance, respectively (Factor 3’s eigenvalue fell below 1 at 0.968). The scree plot likewise outlines a two-factor solution (Supplementary Figure 4). As the two factors were modestly correlated (*r*=.285), an oblique (promax) rotation with Kaiser normalisation was applied, and this pattern of loadings was finally interpreted (Supplementary Table 5).

| BP_ND_^0^ ROIs | Factor | |
| --- | --- | --- |
|  | 1 | 2 |
| Amygdala | **0.469** | 0.185 |
| Caudate | 0.194 | **0.651** |
| Hippocampus | **0.562** | 0.098 |
| LC | **0.338** | 0.232 |
| NAcc | -0.069 | **0.510** |
| Putamen | 0.041 | **0.584** |
| SN | **0.777** | -0.413 |
| Thalamus | **0.723** | 0.100 |
| VTA | **0.740** | 0.069 |

**Supplementary Table 5. Pattern matrix extracted using PAF method.** Rotation method was promax with Kaiser normalisation, and the rotation was converged in 3 iterations. Abbreviations: LC=locus coeruleus; NAcc=nucleus accumbens; SN=substantia nigra; VTA=ventral tegmental area.

**6. Supplementary Result 1: Correlation Between ROI Volumetric Measures and D2/D3 Receptor Availability**

**(A) Age-controlled volume-BP_ND_^0^ correlations**

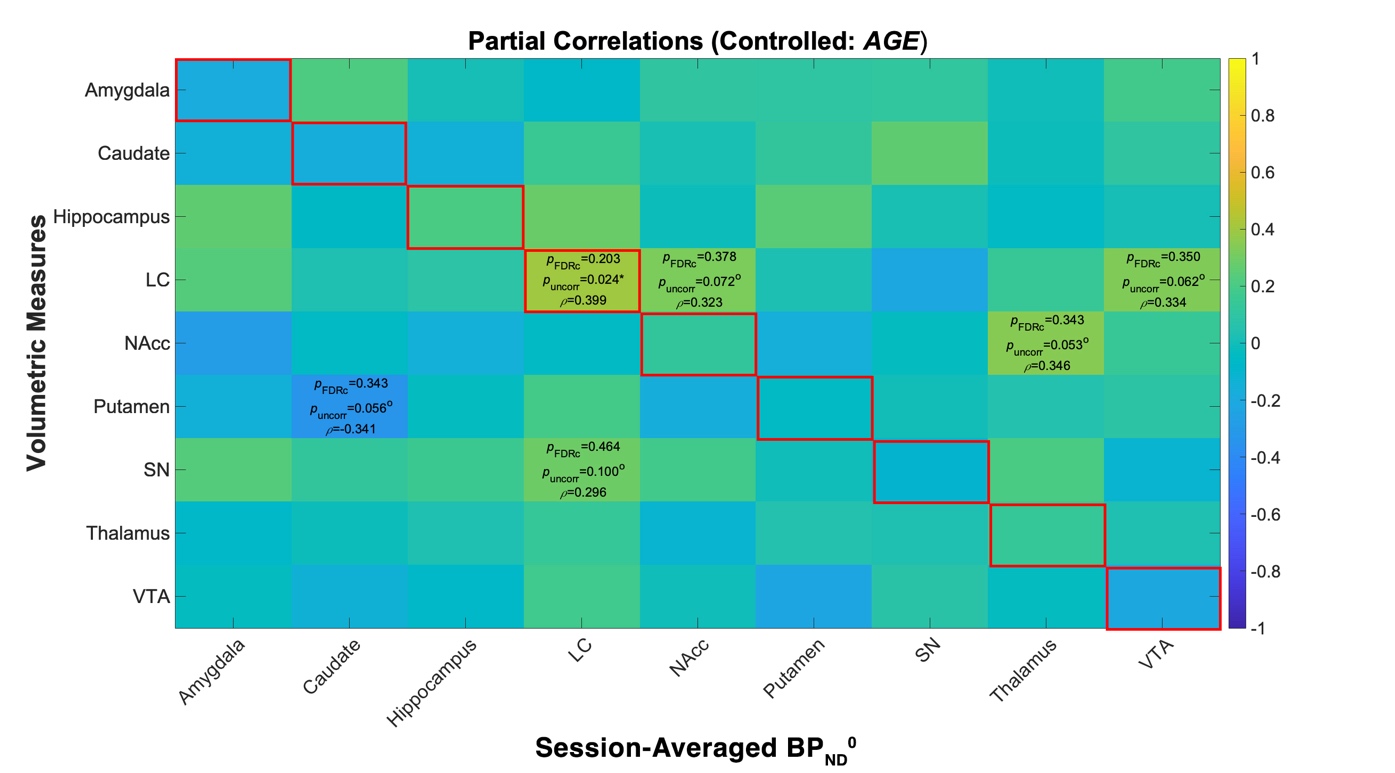

**(B) Age-*un*controlled volume-BP_ND_^0^ correlations**

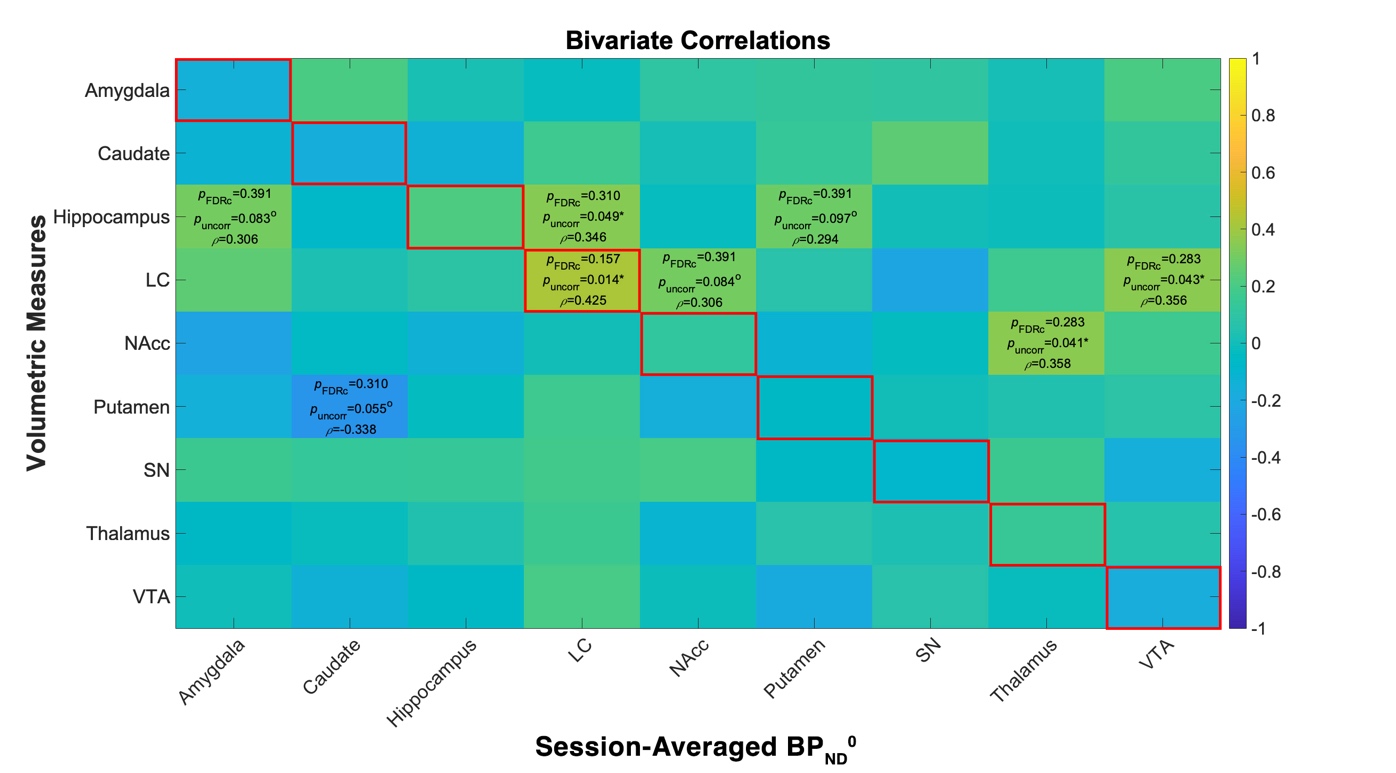

**Supplementary Figure 5. Correlation between volume and BP_ND_^0^ in each and across ROIs in age-controlled and age-uncontrolled nonparametric (Spearman’s Rho) correlational analyses.** **(A)** Age-controlled Spearman correlation matrix. Each cell shows the partial Spearman’s *ρ* between the volume of the ROI on the y-axis and the session-averaged BP_ND_^0^ of the ROI on the x-axis after regressing out age. **(B)** Age-uncontrolled (zero-order) Spearman correlation matrix for the same measures. Colour bars indicate correlation magnitude (blue = negative, yellow = positive). Numbers inside cells give *ρ*, the uncorrected two-tailed (uncorr), and the false-discovery-rate-adjusted (FDRc) p-values. Diagonal cells outlined in red denote volume-BP relationship in a same ROI.

Across all ROIs, none displayed a statistically significant volume-to-BP_ND_^0^ association once we applied the stringent family-wise correction for multiple comparisons (Supplementary Figure 5). Only a small subset of zero-order correlations, as well as age-controlled partial correlations, achieved nominal significance, and these findings did not survive correction. Taken together, these outcomes imply that, within this older cohort sample, inter-individual variability in D2/D3 receptor availability cannot be captured by a simple linear scaling with regional volume. In other words, having a larger caudate, hippocampus, or VTA does not reliably predict higher or lower BP_ND_^0^.

This apparent disjunction is most plausibly rooted in the multifaceted nature of cerebral ageing (Raz and Rodrigue 2006). Structural volumetric loss, microstructural compromise, and dopaminergic receptor decline are each influenced by partially independent biological pathways and can proceed at distinct rates across the brain (Seaman et al. 2019). For instance, vascular and neuroinflammatory factors may drive cortical atrophy without directly altering subcortical D2/D3 receptor expression, whereas dopaminergic denervation may advance in striatum despite preserved grey-matter volume. Such heterogeneity dilutes any straightforward volume-BP relationship at the group level. Taken together, these findings indicate that structural deterioration, microstructural changes, and dopaminergic receptor decline follow partially independent trajectories, so regional volume may be a poor proxy for dopaminergic receptor integrity in healthy older adults. Given these considerations, the isolated nominal findings should be interpreted cautiously.

**6. Supplementary Result 2: Recognition Memory test performance in all trials**

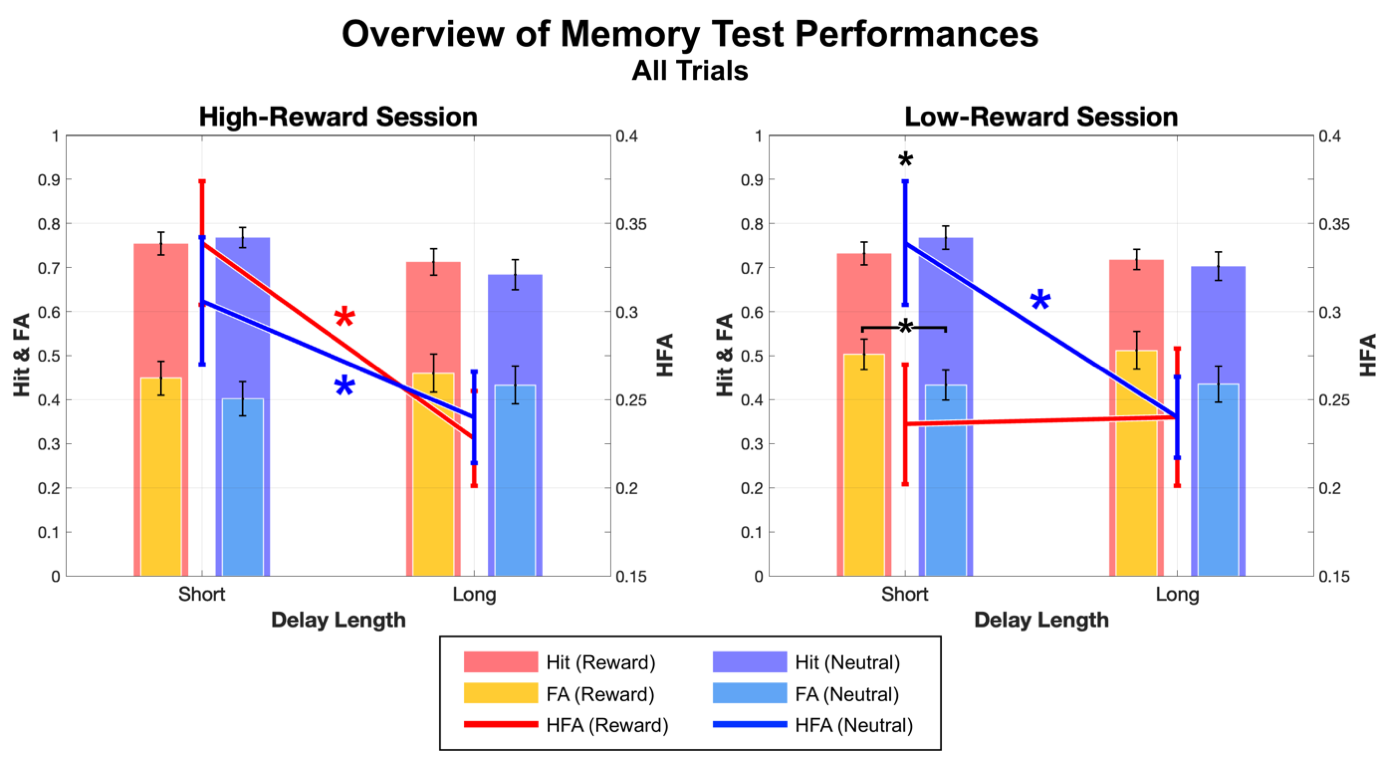

**Supplementary Figure 6. Overview of Recognition Memory test performance in all trials.** This figure compares the compound score of hit rate and FA ($\boldsymbol{Hit -False Alarm}$), i.e. HFA (line graphs), as well as hit rate and FA (nested bar graphs) across two memory delay lengths, short and long, for reward-associated (warm colours) and neutral scenes (cool colours) within each reward context for all trials. The left plot depicts high-reward sessions, and the right plot represents low-reward sessions, illustrating a distinct interaction between the reward context and delay length on overall memory performance. Whiskers around the marker in this plot represents standard error and an asterisk (*) represents statistical significance of p<.05. Again, the memory performance of low-reward sessions’ reward-associated scenes show an atypical pattern across temporal delays, and this pattern seems to be driven by the significant difference between FA for short-delay memory test in reward-associated and neutral scenes.

In the investigation of the effects of reward on subsequent memory retention in healthy older adults, the analysis on various memory performance measures including all trials (trials rated either ‘sure’ or ‘not sure’), revealed similar patterns to those from the analysis that included only high-confidence trials. In a three-factor repeated measures ANOVA (factors: Reward [Reward/Neutral], Delay Length [Short/Long], and Session Context [High-Reward/Low-Reward]), the primary memory performance outcome, HFA, indicated a significant main effect of delay length, with superior memory performance observed in the immediate (short delay; max. 15m after encoding has concluded) post-task recognition memory tests compared to those administered after a 24-hour delay (long delay), *F*(1,32)=9.161, *p*=0.005 (Figure 3F). No significant main effects of Reward or Session Context were found on HFA. A significant three-way interaction between Session Context, Reward, and Delay Length was observed in HFA, *F*(1,32)=8.825, *p*=0.006, *η*²=0.216. Follow-up analyses within the low-reward context showed that HFA for neutral scenes significantly declined from short to long delay, whereas no significant change was observed across delays for reward-associated scenes.

A separate follow-up repeated measures ANOVA analyses on hit rates showed a significant main effect of Delay Length, *F*(1,32)=4.646, *p*=0.039, with higher hit rates at the short delay compared to the long delay. No additional main effects or interactions on hit rates were found in this analyis. For FA, there was significant main effect of Reward, *F*(1,32)=6.098, *p*=0.019, indicating that reward contingency adversely affected FA. Interestingly, this disadvantage of reward salience on FA was most prominent in short-delay tests in low motivational context, *F*(1,32)=6.702, *p*=0.014, *η*²=0.173. Likewise, no other main effects or interaction effect among any factors was found statistically significant.

**6. Supplementary Result 3: Session-specific correlation of BP_ND_^0^ and memory performance indices**

(A) **Non-parametric Partial Correlation Between BP_ND_^0^ and HFA**

**(A-1) High-reward session**

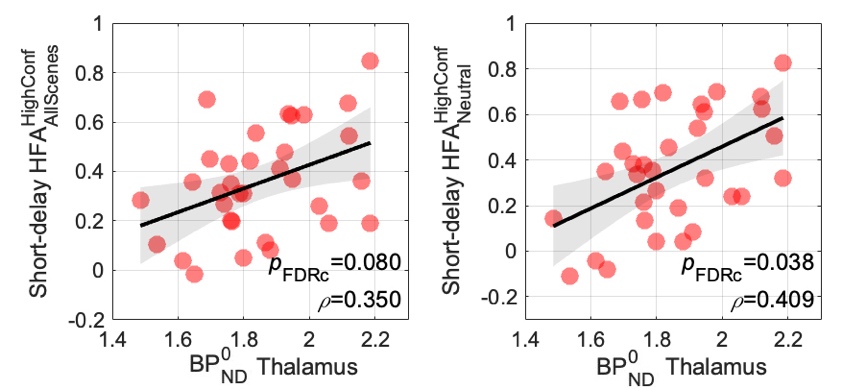

**(A-2) Low-reward session**

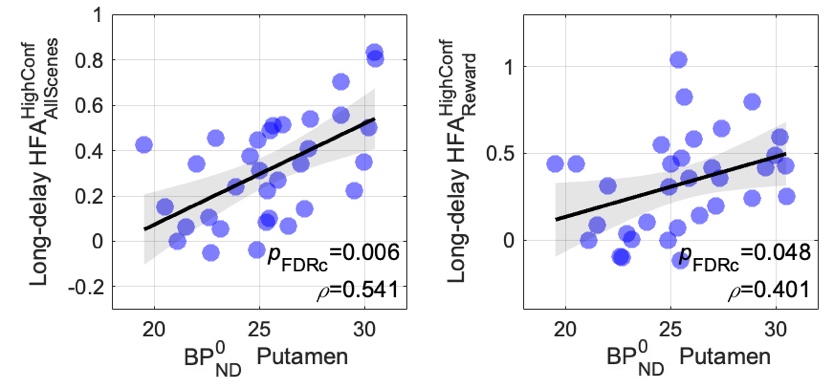

(B) **Non-parametric Partial Correlation Between BP_ND_^0^ and Hit rates**

**(B-1) High-reward session**

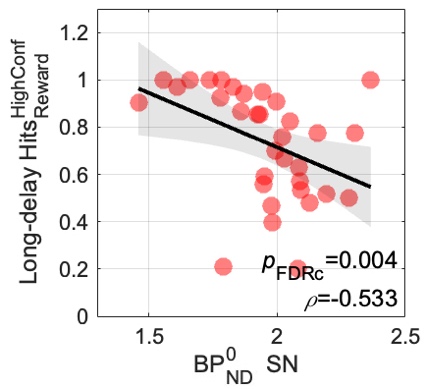

**(B-2) Low-reward session**

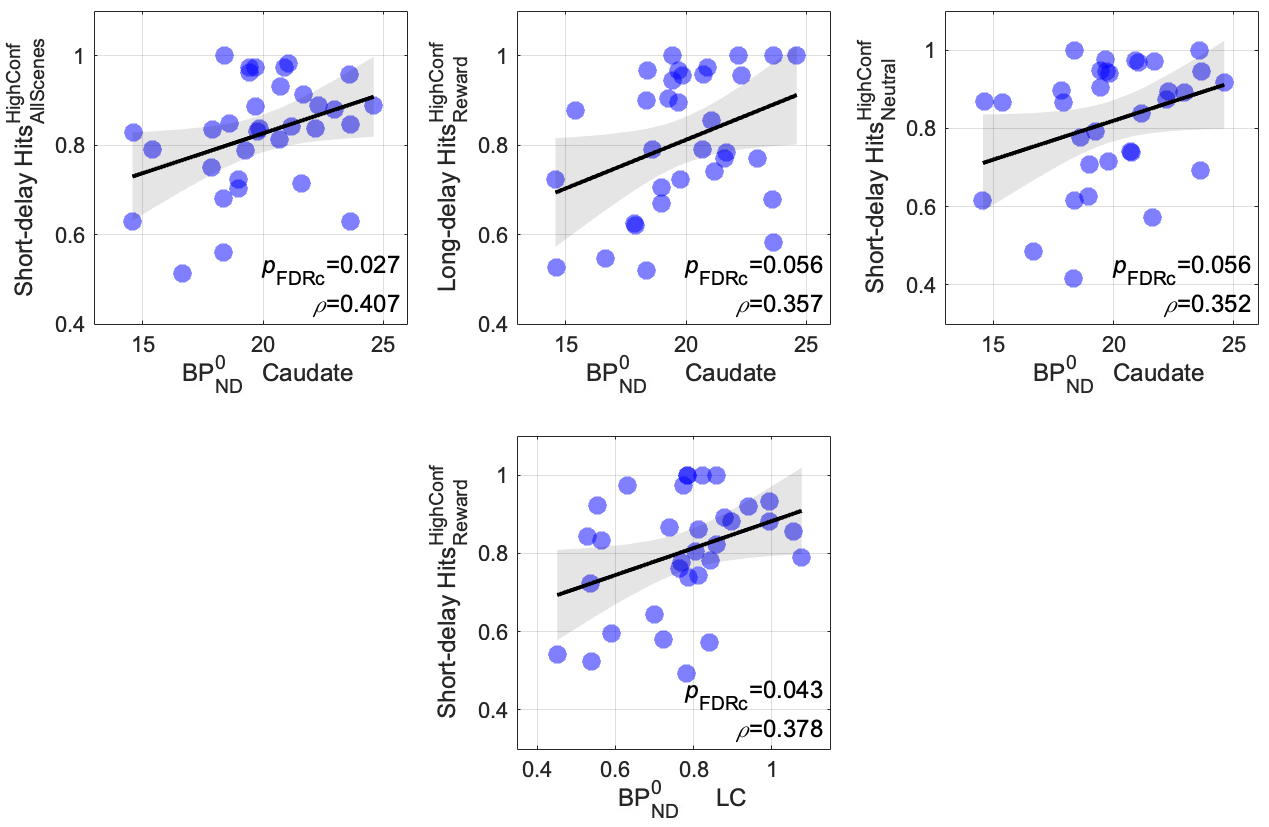

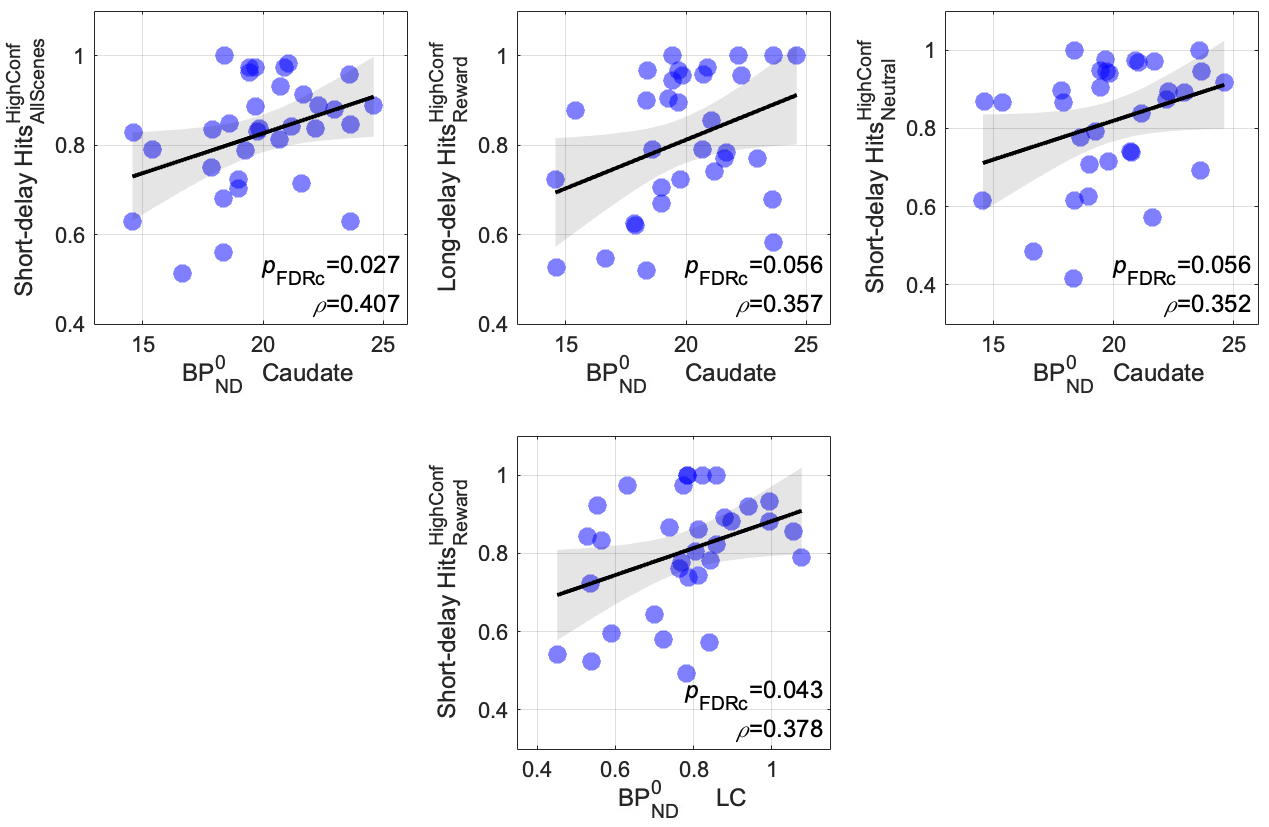

(C) **Non-parametric Partial Correlation Between BP_ND_^0^ and FA**

**(C-1) High-reward session**

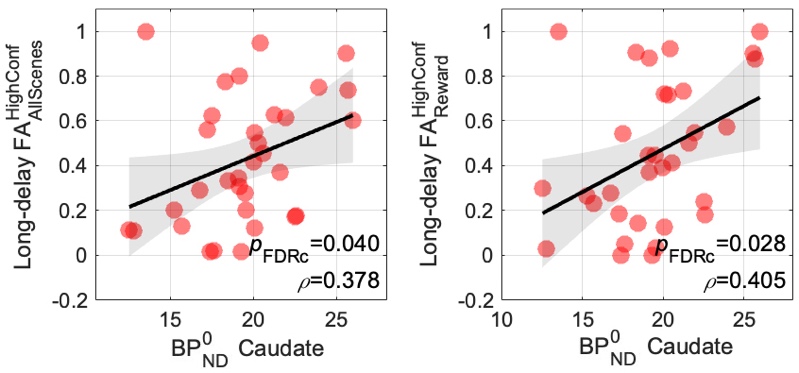

**(C-2) Low-reward session**

*No significant correlation found*

**Supplementary Figure 7. Age-controlled non-parametric correlation between baseline D2/D3 receptor availability and memory performance indices within different sessions.** The above scatterplot figures illustrate the results of correlational analyses between BP_ND_^0^ measured in each ROI and memory performance indices, i.e. **(A) HFA**, **(B) hit rates**, and **(C) FA**. The indices were calculated by only including high-confidence trials of subsequent incidental memory tests assessed at two delay intervals, short- and long-delay (~15 minutes and 24 hours after encoding, respectively). Each dot represents a single data point, and the black solid lines indicate linear fits, with shaded grey areas denoting the standard error. Within each plot, the Spearman's rho (*ρ*) and FDR-corrected p-value (*p*_FDRc_) are displayed in the bottom right corner.

To examine context-dependent relationships between regional D2/D3 receptor availability and memory performance, we performed age-controlled, non-parametric partial correlations between BP_ND_^0^ and memory indices, high-confidence HFA, hit rates, and FA, separately for the high-reward and low-reward sessions and for short- and long-delay tests. In the high-reward session, BP_ND_^0^ in the thalamus showed a trend-level positive association with short-delay HFA across all scenes, *ρ*(32)=0.350, *p*_uncorr_=0.049, *p*_FDRc_=0.080, and a significant positive correlation with that of neutral scenes, *ρ*(32)=0.409, *p*_uncorr_=0.020, *p*_FDRc_=0.038. In the low-reward session, subjects who showed high BP_ND_^0^ in the putamen tend to show better discriminability for all scenes in long-delay recognition tests, *ρ*(32)=0.541, *p*_uncorr_=0.001, *p*_FDRc_=0.006, and also for reward-associated scenes, *ρ*(32)=0.401, *p*_uncorr_=0.023, *p*_FDRc_=0.048.

In the analysis with hit rates, high SN BP_ND_^0^ in the high-reward session predicted *lower* long-delay hit rates for reward-associated scenes in subjects, *ρ*(32)=-0.533, *p*_uncorr_=0.002, *p*_FDRc_=0.004. On the other hand, low-reward BP_ND_^0^ in the caudate showed a significant positive relationship with short-delay hit rates across all scenes, *ρ*(32)=0.407, *p*_uncorr_=0.021, *p*_FDRc_=0.027, and trend-level positive associations with hit rates for reward scenes at long delay, *ρ*(32)=0.357, *p*_uncorr_=0.045, *p*_FDRc_=0.056, and for neutral scenes at short delay, *ρ*(32)=0.352, *p*_uncorr_=0.048, *p*_FDRc_=0.056.

In addition, interestingly, BP_ND_^0^ in the LC was positively correlated with short-delay hit rates for reward-associated scenes in the low-reward session, *ρ*(32)=0.378, *p*_uncorr_=0.033, *p*_FDRc_=0.043. Higher D2/D3 receptor availability in the LC predicted higher short-delay hit rates for reward-associated scenes (Supplementary Figure 7B-2). Importantly, this relationship was not present in the cross-session dataset, for HFA rates, or for long-delay memory test performance. This highly specific pattern highlights a subtle and context-dependent role of LC as a part of dopaminergic system in memory. Despite BP_ND_^0^ index being a stable, trait-like measure of D2/D3 receptor availability, its functional association with immediate reward memory was exclusively observed in session-specific analyses, specifically only in low-motivational context. This result likely signifies a system better equipped to mediate attentional allocation and goal-directed behaviour especially when overall motivational salience is low and focus becomes more important.

The LC-noradrenaline (NA) system is critical for enhancing the signal-to-noise ratio in cortical processing, focusing attention on prioritised, goal-relevant stimuli, and modulating neural gain during encoding of salient information (Mather et al. 2016; Sara 2009). For reward-associated scenes, individuals with a more robust LC function (higher D2/D3 receptor availability) may exhibit superior attentional modulation and deeper encoding of these motivationally relevant items. The fact that D2/D3 receptor availability in the LC correlated only with hit rates for reward-associated scenes and not with HFA or FA implies that LC’s influence may be selective to enhancement of goal-relevant stimuli without improving general sharpness of memory. Similar to the correlation found in the thalamic D2/D3 receptor availability and short-delay memory test performance, while NA, the LC's primary output, is crucial for memory consolidation (Dahl et al. 2023; Gibbs, Hutchinson, and Summers 2010), long-delay memory likely relies on more complex, dynamic, and distributed consolidation processes that extend beyond what a baseline receptor measure singularly predicts.

Finally, FA in the high-reward session were uniquely predicted by BP_ND_^0^ in the caudate, which correlated positively with long-delay FA across all scenes, *ρ*(32)=0.378, *p*_uncorr_=0.033, *p*_FDRc_=0.040, and for reward-associated scenes, *ρ*(32)=0.405, *p*_uncorr_=0.022, *p*_FDRc_=0.028. No correlations between BP_ND_^0^ and FA survived correction in the low-reward session.

Karlsson, Max, Cheng Zhang, Loren Méar, Wen Zhong, Andreas Digre, Borbala Katona, Evelina Sjöstedt, Lynn Butler, Jacob Odeberg, Philip Dusart, Fredrik Edfors, Per Oksvold, Kalle Von Feilitzen, Martin Zwahlen, Muhammad Arif, Ozlem Altay, Xiangyu Li, Mehment Ozcan, Adil Mardinoglu, Linn Fagerberg, Jan Mulder, Yonglun Luo, Fredrik Ponten, Mathias Uhlén, and Cecilia Lindskog. 2021. ‘A Single–Cell Type Transcriptomics Map of Human Tissues’. *SCIENCE ADVANCES*.

Lammertsma, Adriaan A., and Susan P. Hume. 1996. ‘Simplified Reference Tissue Model for PET Receptor Studies’. *NeuroImage* 4(3):153–58. doi:10.1006/nimg.1996.0066.

Little, Roderick J. A. 1988. ‘A Test of Missing Completely at Random for Multivariate Data with Missing Values’. *Journal of the American Statistical Association* 83(404):1198–1202. doi:10.1080/01621459.1988.10478722.

Mather, Mara, David Clewett, Michiko Sakaki, and Carolyn W. Harley. 2016. ‘Norepinephrine Ignites Local Hotspots of Neuronal Excitation: How Arousal Amplifies Selectivity in Perception and Memory’. *Behavioral and Brain Sciences* 39:e200. doi:10.1017/S0140525X15000667.

McCombe, Niamh, Shuo Liu, Xuemei Ding, Girijesh Prasad, Magda Bucholc, David P. Finn, Stephen Todd, Paula L. McClean, and KongFatt Wong-Lin. 2022. ‘Practical Strategies for Extreme Missing Data Imputation in Dementia Diagnosis’. *IEEE Journal of Biomedical and Health Informatics* 26(2):818–27. doi:10.1109/JBHI.2021.3098511.

Pauli, Wolfgang M., Amanda N. Nili, and J. Michael Tyszka. 2018. ‘A High-Resolution Probabilistic in Vivo Atlas of Human Subcortical Brain Nuclei’. *Scientific Data* 5(1):180063. doi:10.1038/sdata.2018.63.

Raz, Naftali, and Karen M. Rodrigue. 2006. ‘Differential Aging of the Brain: Patterns, Cognitive Correlates and Modifiers’. *Neuroscience & Biobehavioral Reviews* 30(6):730–48. doi:10.1016/j.neubiorev.2006.07.001.

Sara, Susan J. 2009. ‘The Locus Coeruleus and Noradrenergic Modulation of Cognition’. *Nature Reviews Neuroscience* 10(3):211–23. doi:10.1038/nrn2573.

Schafer, Joseph L., and John W. Graham. 2002. ‘Missing Data: Our View of the State of the Art.’ *Psychological Methods* 7(2):147–77. doi:10.1037/1082-989X.7.2.147.

Seaman, Kendra L., Christopher T. Smith, Eric J. Juarez, Linh C. Dang, Jaime J. Castrellon, Leah L. Burgess, M. Danica San Juan, Paul M. Kundzicz, Ronald L. Cowan, David H. Zald, and Gregory R. Samanez‐Larkin. 2019. ‘Differential Regional Decline in Dopamine Receptor Availability across Adulthood: Linear and Nonlinear Effects of Age’. *Human Brain Mapping* 40(10):3125–38. doi:10.1002/hbm.24585.

Shishegar, Rosita, Timothy Cox, David Rolls, Pierrick Bourgeat, Vincent Doré, Fiona Lamb, Joanne Robertson, Simon M. Laws, Tenielle Porter, Jurgen Fripp, Duygu Tosun, Paul Maruff, Greg Savage, Christopher C. Rowe, Colin L. Masters, Michael W. Weiner, Victor L. Villemagne, and Samantha C. Burnham. 2021. ‘Using Imputation to Provide Harmonized Longitudinal Measures of Cognition across AIBL and ADNI’. *Scientific Reports* 11(1):23788. doi:10.1038/s41598-021-02827-6.

Sjöstedt, Evelina, Wen Zhong, Linn Fagerberg, Max Karlsson, Nicholas Mitsios, Csaba Adori, Per Oksvold, Fredrik Edfors, Agnieszka Limiszewska, Feria Hikmet, Jinrong Huang, Yutao Du, Lin Lin, Zhanying Dong, Ling Yang, Xin Liu, Hui Jiang, Xun Xu, Jian Wang, Huanming Yang, Lars Bolund, Adil Mardinoglu, Cheng Zhang, Kalle Von Feilitzen, Cecilia Lindskog, Fredrik Pontén, Yonglun Luo, Tomas Hökfelt, Mathias Uhlén, and Jan Mulder. 2020. ‘An Atlas of the Protein-Coding Genes in the Human, Pig, and Mouse Brain’. *Science* 367(6482):eaay5947. doi:10.1126/science.aay5947.

Slifstein, Mark, and Marc Laruelle. 2000. ‘Effects of Statistical Noise on Graphic Analysis of PET Neuroreceptor Studies’. *THE JOURNAL OF NUCLEAR MEDICINE* 41(12):2083–88.

Thul, Peter J., Lovisa Åkesson, Mikaela Wiking, Diana Mahdessian, Aikaterini Geladaki, Hammou Ait Blal, Tove Alm, Anna Asplund, Lars Björk, Lisa M. Breckels, Anna Bäckström, Frida Danielsson, Linn Fagerberg, Jenny Fall, Laurent Gatto, Christian Gnann, Sophia Hober, Martin Hjelmare, Fredric Johansson, Sunjae Lee, Cecilia Lindskog, Jan Mulder, Claire M. Mulvey, Peter Nilsson, Per Oksvold, Johan Rockberg, Rutger Schutten, Jochen M. Schwenk, Åsa Sivertsson, Evelina Sjöstedt, Marie Skogs, Charlotte Stadler, Devin P. Sullivan, Hanna Tegel, Casper Winsnes, Cheng Zhang, Martin Zwahlen, Adil Mardinoglu, Fredrik Pontén, Kalle Von Feilitzen, Kathryn S. Lilley, Mathias Uhlén, and Emma Lundberg. 2017. ‘A Subcellular Map of the Human Proteome’. *Science* 356(6340):eaal3321. doi:10.1126/science.aal3321.

Uhlén, Mathias, Linn Fagerberg, Björn M. Hallström, Cecilia Lindskog, Per Oksvold, Adil Mardinoglu, Åsa Sivertsson, Caroline Kampf, Evelina Sjöstedt, Anna Asplund, IngMarie Olsson, Karolina Edlund, Emma Lundberg, Sanjay Navani, Cristina Al-Khalili Szigyarto, Jacob Odeberg, Dijana Djureinovic, Jenny Ottosson Takanen, Sophia Hober, Tove Alm, Per-Henrik Edqvist, Holger Berling, Hanna Tegel, Jan Mulder, Johan Rockberg, Peter Nilsson, Jochen M. Schwenk, Marica Hamsten, Kalle Von Feilitzen, Mattias Forsberg, Lukas Persson, Fredric Johansson, Martin Zwahlen, Gunnar Von Heijne, Jens Nielsen, and Fredrik Pontén. 2015. ‘Tissue-Based Map of the Human Proteome’. *Science* 347(6220):1260419. doi:10.1126/science.1260419.

Uhlén, Mathias, Max J. Karlsson, Andreas Hober, Anne-Sophie Svensson, Julia Scheffel, David Kotol, Wen Zhong, Abdellah Tebani, Linnéa Strandberg, Fredrik Edfors, Evelina Sjöstedt, Jan Mulder, Adil Mardinoglu, Anna Berling, Siri Ekblad, Melanie Dannemeyer, Sara Kanje, Johan Rockberg, Magnus Lundqvist, Magdalena Malm, Anna-Luisa Volk, Peter Nilsson, Anna Månberg, Tea Dodig-Crnkovic, Elisa Pin, Martin Zwahlen, Per Oksvold, Kalle Von Feilitzen, Ragna S. Häussler, Mun-Gwan Hong, Cecilia Lindskog, Fredrik Ponten, Borbala Katona, Jimmy Vuu, Emil Lindström, Jens Nielsen, Jonathan Robinson, Burcu Ayoglu, Diana Mahdessian, Devin Sullivan, Peter Thul, Frida Danielsson, Charlotte Stadler, Emma Lundberg, Göran Bergström, Anders Gummesson, Bjørn G. Voldborg, Hanna Tegel, Sophia Hober, Björn Forsström, Jochen M. Schwenk, Linn Fagerberg, and Åsa Sivertsson. 2019. ‘The Human Secretome’. *Science Signaling* 12(609):eaaz0274. doi:10.1126/scisignal.aaz0274.

Uhlen, Mathias, Max J. Karlsson, Wen Zhong, Abdellah Tebani, Christian Pou, Jaromir Mikes, Tadepally Lakshmikanth, Björn Forsström, Fredrik Edfors, Jacob Odeberg, Adil Mardinoglu, Cheng Zhang, Kalle Von Feilitzen, Jan Mulder, Evelina Sjöstedt, Andreas Hober, Per Oksvold, Martin Zwahlen, Fredrik Ponten, Cecilia Lindskog, Åsa Sivertsson, Linn Fagerberg, and Petter Brodin. 2019. ‘A Genome-Wide Transcriptomic Analysis of Protein-Coding Genes in Human Blood Cells’. *Science* 366(6472):eaax9198. doi:10.1126/science.aax9198.

Uhlen, Mathias, Cheng Zhang, Sunjae Lee, Evelina Sjöstedt, Linn Fagerberg, Gholamreza Bidkhori, Rui Benfeitas, Muhammad Arif, Zhengtao Liu, Fredrik Edfors, Kemal Sanli, Kalle Von Feilitzen, Per Oksvold, Emma Lundberg, Sophia Hober, Peter Nilsson, Johanna Mattsson, Jochen M. Schwenk, Hans Brunnström, Bengt Glimelius, Tobias Sjöblom, Per-Henrik Edqvist, Dijana Djureinovic, Patrick Micke, Cecilia Lindskog, Adil Mardinoglu, and Fredrik Ponten. 2017. ‘A Pathology Atlas of the Human Cancer Transcriptome’. *Science* 357(6352):eaan2507. doi:10.1126/science.aan2507.

Vernaleken, Ingo, Lisa Peters, Mardjan Raptis, Robert Lin, Hans-Georg Buchholz, Yun Zhou, Oliver Winz, Frank Rösch, Peter Bartenstein, Dean F. Wong, Wolfgang M. Scháfer, and Gerhard Gründer. 2011. ‘The Applicability of SRTM in [^18^ F]Fallypride PET Investigations: Impact of Scan Durations’. *Journal of Cerebral Blood Flow & Metabolism* 31(9):1958–66. doi:10.1038/jcbfm.2011.73.
